# In vivo elastography of the human retina using light-evoked intrinsic actuation

**DOI:** 10.64898/2026.05.01.722017

**Authors:** Teng Liu, Huakun Li, Vimal Prabhu Pandiyan, Keyu Chen, Palash Bharadwaj, Benjamin J. Wendel, Debarshi Mustafi, Jennifer R. Chao, Tong Ling, Ramkumar Sabesan

## Abstract

The biomechanical properties of the retina govern its function, structural integrity, and susceptibility to disease, yet remain difficult to measure in vivo due to the lack of safe, spatially localized mechanical actuation. Here, we introduce a framework for probing retinal biomechanics in the living human eye by leveraging intrinsic optical actuation driven by phototransduction. Using phase-resolved optical coherence tomography with a local phase-referencing approach, we resolved signed, nanometer-scale displacements of the major outer retinal interfaces evoked by light. The resulting deformation field, originating in the photoreceptor outer segment, was distributed across retinal compartments in an eccentricity-dependent manner, with efficient axial transfer in the fovea and attenuated propagation in the parafovea. A hybrid analytical and finite-element framework was developed that retrieved the biomechanical properties of the retinal compartments based on their coordinated deformation and the anatomical variation in retinal structure versus eccentricity. In retinitis pigmentosa, the paradigm enabled the detection of light-evoked deformation in the transition zone despite the loss of native lamination, enabling a functional readout of the vulnerable photoreceptors at the leading edge of degeneration. Together, these results establish intrinsic optical stimulation as a basis for in vivo retinal elastography and enable the non-invasive, quantitative imaging of retinal biomechanics and function in the living human retina.

## Introduction

Biomechanical properties of tissues arise from the interplay between anatomical architecture, cellular composition, and material properties, and are fundamental to structural integrity, tissue homeostasis, and physiological function^1–3^. Altered biomechanics have been implicated in development, aging and disease^4–6^. Traditional approaches to assess tissue mechanics rely on ex vivo preparations or externally applied forces^7–10^, that are limited in their ability to replicate the native physiological environment, including boundary conditions and geometry^11^, that govern mechanical and physiological behavior in vivo^8,9^. In some tissues, however, endogenous physiological processes can serve as intrinsic sources of mechanical actuation, enabling non-invasive and functionally relevant assessment of biomechanics^12^. Examples include the liver and brain, where respiration and hemodynamics generate internal forces deforming tissue which can then be measured using imaging-based elastography approaches^12–14^.

The retina presents a unique opportunity to explore the use of endogenous forces for mapping biomechanical properties. Major diseases including age-related macular degeneration (AMD), hereditary retinal dystrophies, vitreoretinal interface disorders and glaucoma involve changes in tissue structure and organization that are likely accompanied, or even preceded by alterations in mechanical properties^10,15–18^. Understanding retinal biomechanical properties is also relevant for surgical handling and the design of subretinal therapies and implants^19–22^. While the structural organization of the retina - its laminar architecture, cellular composition and microanatomy - is well characterized by anatomical studies^23–25^ and high-resolution imaging^26^, its mechanical properties remain far less understood. Biomechanical insight has largely been derived from ex vivo tissue preparations, such as the tensile testing of donor tissue, atomic force microscopy (AFM) of isolated human inner limiting membrane (ILM), whole-globe inflation, and vitreoretinal peel assays^10^. These ex vivo approaches are inherently limited by post-mortem processing and the resultant disruption of native tissue environment. Quantitative in vivo assessment of retinal biomechanics in a living eye remains challenging given that its thin, multilayered structure is anatomically situated at the back of the eye, rendering it largely inaccessible for administering a controlled, external perturbation with high spatial specificity.

Current ocular biomechanical assays have shown promise when applied in vivo to the cornea^27,28^, lens^29^ and sclera^27^, but remain limited in their application to the retina. For instance, Brillouin microscopy^30,31^ takes advantage of the frequency shifts of light waves passing through heterogeneous tissue, but this approach is limited to the anterior eye and is yet to be applied in vivo in the retina. Optical coherence elastography (OCE) has been used to study the retinal tissue in animal models^32^, but relies on external force generation (e.g.: acoustic radiation force, shaker-based mechanical excitation^33,34^) that may exceed ocular safety limits in human studies of the retina^35^. Overall, these limitations motivate the development of approaches that exploit endogenous physiological actuation with high spatial specificity for in vivo interrogation of retinal biomechanics.

Here, we sought to take advantage of an intrinsic phenomenon of the retina – phototransduction – whereby a light stimulus is converted to electrical activity through a well-characterized cascade of amplification steps^36^. The transduction of light initiates in the photoreceptor outer segment and is accompanied by electrical and osmotic changes that result in nanometer-scale axial deformations^37,38^. These displacements can be detected with phase-resolved optical coherence tomography (OCT)^38–40^ and form the basis of optoretinography (ORG)^40,41,38,42–44^. We asked whether such light-evoked deformations could be linked quantitatively to cellular and tissue biomechanics. A key challenge, however, is that these deformations are typically measured as changes in optical path length between pairs of reflective interfaces, each associated with distinct neighboring compartments and boundary conditions. As a result, the measured signals reflect relative separation between layers and cannot distinguish whether either or both interfaces have moved, or by how much. This precludes the assignment of motion to individual layers and limiting quantitative interpretation of these deformations in terms of underlying mechanical properties.

To address this limitation, we developed a lateral reference-based approach using localized stimulation. This local referencing paradigm enables the recovery of layer-specific displacements in response to light stimulation across the entire thickness of the retina and, when combined with adaptive optics (AO), at the cellular scale. In conjunction with a hybrid analytical spring series and finite element model, this approach enables inference of the mechanical properties of distinct outer retinal compartments in regions with markedly different anatomical specialization, establishing a framework for the in vivo assessment of retinal biomechanics using intrinsic optical actuation. Finally, we apply this framework in patients with inherited retinal disease and show alterations consistent with early retinal remodeling.

## Results

### Deformation in outer retinal compartments in response to light

Light-evoked deformations were measured using a custom-built line-scan OCT system^45^ from the outer retina (see Protocol 1 in Methods). Figure 1A shows a representative cross-sectional (B-scan) of the human retina, where several hyperreflective outer retinal interfaces, including the external limiting membrane (ELM), inner segment/outer segment junction (ISOS), cone outer segment tip (COST), and retinal pigment epithelium (RPE) are apparent. To quantify the outer retinal deformations following visual stimulation, temporal changes in optical path length (ΔOPL) between five pairs of retinal interfaces were computed (see colored arrows in Fig. 1B). These include the two boundaries encasing the cone inner segment (IS; ELM–ISOS) and the outer segment (OS; ISOS–COST), as well as between RPE and the other major interfaces (COST–RPE, ISOS–RPE, and ELM–RPE). In response to a green flash of 14.1 × 10^6^ photons⁄μm^2^ on the retina, the cone OS (yellow traces in Fig. 1C, ISOS-COST) expanded rapidly after stimulus onset, approaching a sustained plateau of ∼900 nm at 0.5°, whereas at 5° the response saturated at a smaller amplitude (∼400 nm), similar in trend and magnitude vs. eccentricity as previous reports^45^. Over the same interval, ΔOPL in the supracone space (SCS, COST–RPE) decreased, while Δ OPL spanning the cone OS and supracone space (ISOS–RPE) increased. Notably, compartment spanning the ELM to the RPE expanded near the fovea by ∼100 nm in ∼0.4 s. At 5° eccentricity, this compartment remained stationary within the measurements’ noise floor (cyan curves in Fig. 1C). Finally, the inner segments (spanning the ELM to the ISOS junction) exhibited a contraction of ∼80 nm at 0.5° and 60 nm at 5°.

**Figure 1.**
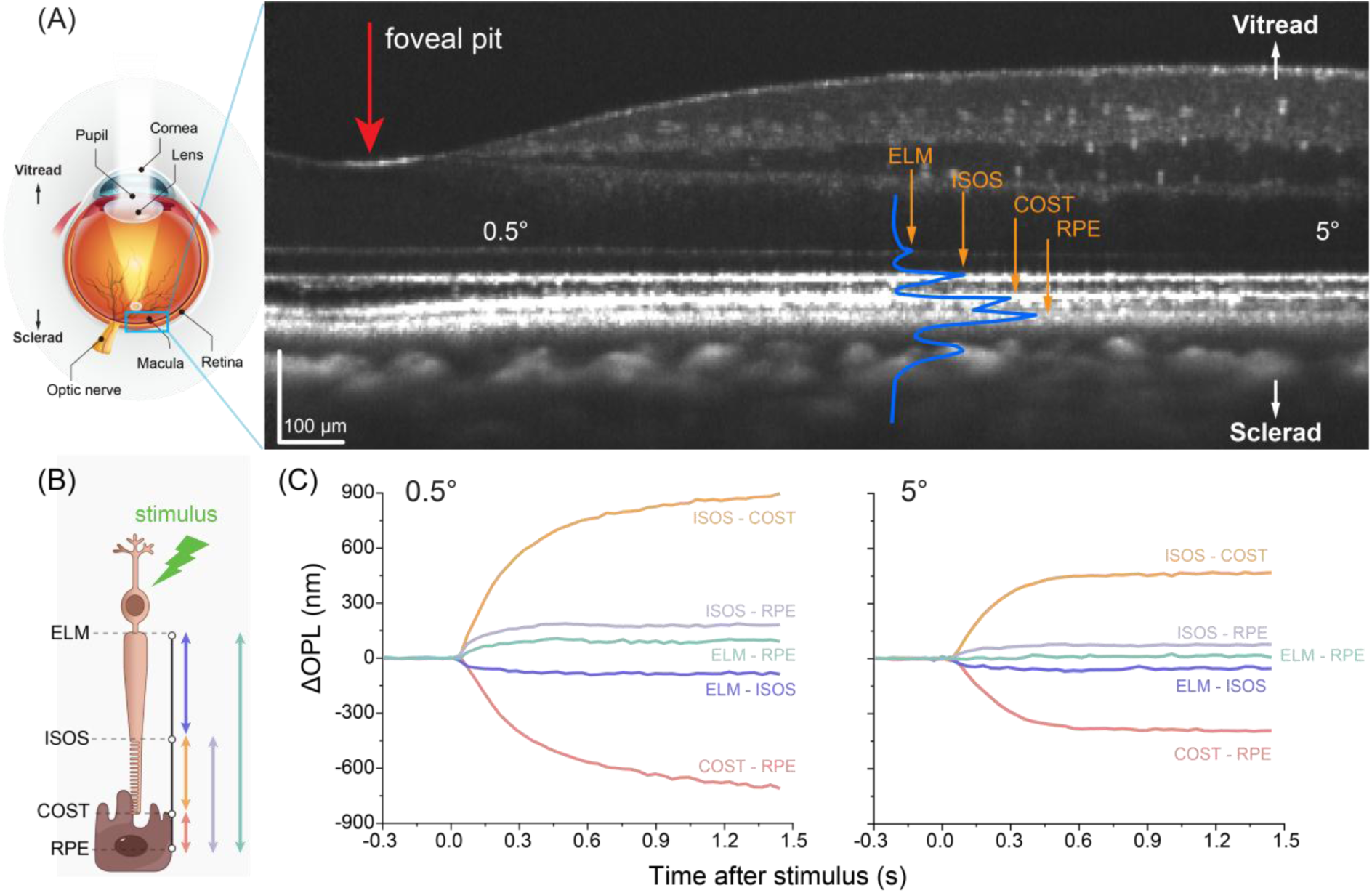
Light-evoked deformations in outer retinal compartments (A) Left: Illustration of the human eye highlighting key anatomical components and the axial direction, with “vitread” and “sclerad” denoting directions toward the vitreous and sclera, respectively. Right: representative retinal cross-section (OCT B-scan) spanning the central retina from the fovea to 5° eccentricity. The blue curve represents the axial intensity profile. Foveal pit is indicated by the red arrow. ELM: external limiting membrane; ISOS: inner-segment/outer-segment junction; COST: cone outer segment tip; RPE: retinal pigment epithelium. (B) Schematic of a cone photoreceptor and the underlying RPE. Colored arrows illustrate layer pairs between which the light-evoked deformations are quantified as changes in optical path length (Δ OPL). (C) Representative time courses of ΔOPL for the five layer-pairs at 0.5° (left panel) and 5° (right panel) in response to a green flash with a photon density of 14.1 × 10^6^ photons⁄μm^2^ on the retina.

### Local phase referencing enables detection of absolute, layer-resolved retinal displacements in response to light

Overall, Fig. 1 shows that light-evoked deformations initiated in the cone OS propagates to neighboring retinal compartments. This indicates that phototransduction provides an intrinsic mechanical drive capable of deforming the layered structure of the outer retina, offering a potential avenue to probe its biomechanics without external mechanical loading. The outer retina comprises multiple compartments with distinct structure and material properties delineated by specific interfaces and their associated boundary constraints. Accordingly, the partitioning of deformation across interfaces reflects the mechanical coupling between compartments. However, the ΔOPL signals, as measured here and in all prior reports of ORG, represent changes in the *relative* separation between two reflective interfaces and therefore quantify the deformation of the compartments they bound rather than the motion of the interfaces themselves. Under this axial phase-referencing paradigm, in which changes in optical phase or OPL are measured between two axially separated retinal interfaces, the motion of the individual layers cannot be determined directly. To resolve this ambiguity, we developed a local phase-referencing approach that resolves the motion of individual outer retinal interfaces and decomposes the observed compartment deformation into layer specific contributions (Fig. 2).

**Figure 2.**
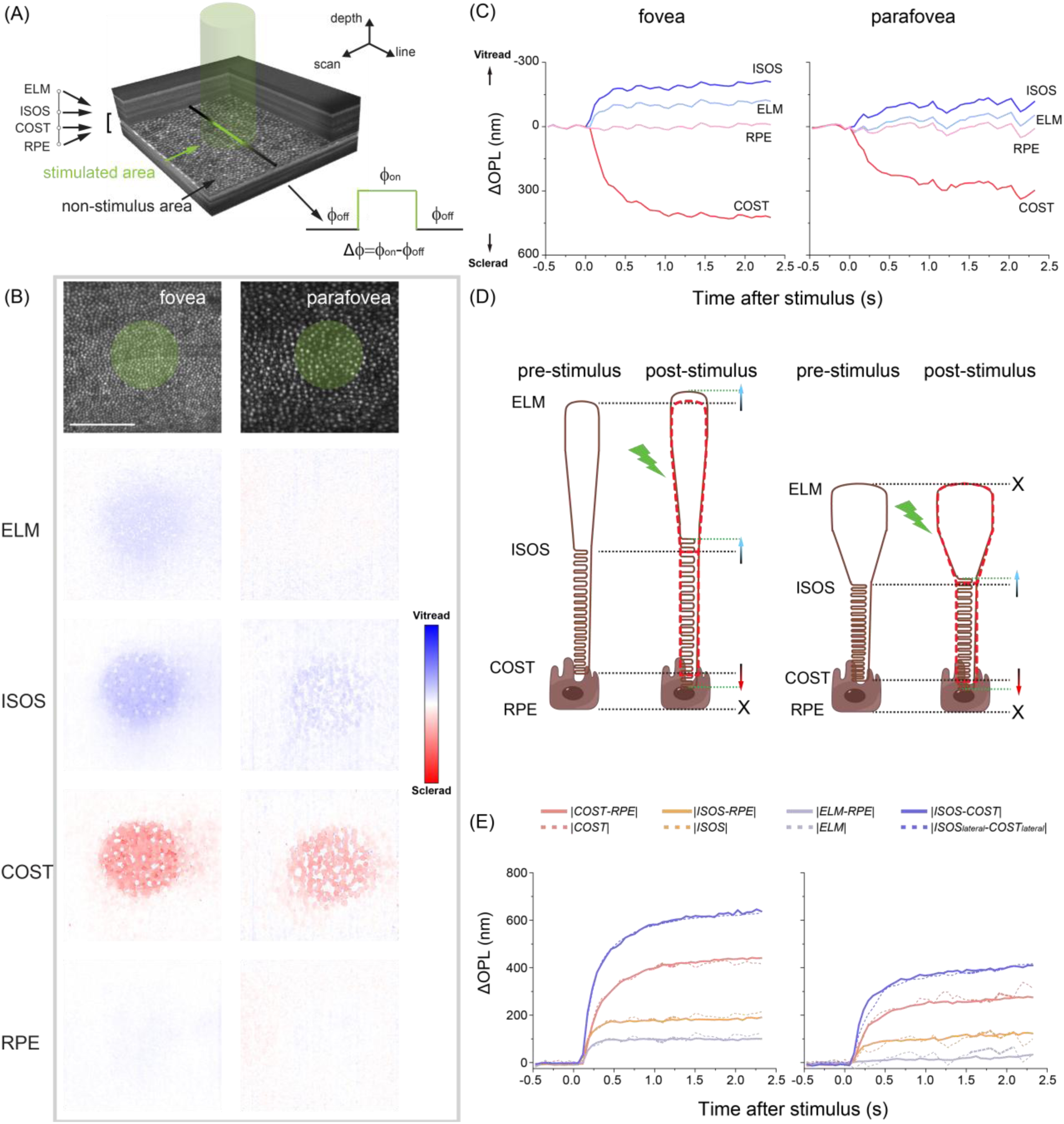
Absolute, layer-resolved displacement of the outer retinal interfaces in response to light (photon density: 8.8 × 10^6^ photons⁄μm^2^), revealed by local phase referencing (A) A spatially localized visual stimulus (green cylinder) was delivered during AO-OCT volume series acquisition. Phase differences were calculated between the stimulated area and the surrounding unstimulated area to yield the absolute displacement of individual retinal interfaces in response to light (also see Supplementary Fig. 1). (B) En face views illustrate the localized stimulation (green disks) overlaid on the cone structural images at the fovea (0.5°) and parafovea (5°), obtained by segmenting the registered AO-OCT volumes at the COST. The local-referenced ΔOPL maps, obtained by averaging the time evolution of light response from 0.4–0.6 s post-stimulus, is shown for the four major outer retinal layers – ELM, ISOS, COST and RPE. Color bar indicates direction of motion. Scale bar: 100 µm. (C) Time evolution of light-evoked displacements for the four layers, plotted as ΔOPL versus time, at the fovea and parafovea. Displacements on the y-axis are expressed as denoting vitread (negative) or sclerad (positive) directions. The oscillatory behavior in the traces reflects pulsations due to heartbeat. (D) Schematic representation of the layer displacements in the fovea (left) and parafovea (right) following visual stimulation. ‘X’ denotes no measurable movement, and red/blue arrows indicate vitread/scleral movement. The red dashed outline in the post-stimulus schematic is presented as a visual guide for the original pre-stimulus locations for the interfaces. (E) Comparison between unsigned axial-referenced ΔOPL traces calculated between various retinal layer pairs (solid curves) and unsigned lateral-referenced ΔOPL traces (dashed curves) at the fovea (left) and parafovea (right). The dashed blue curve was obtained by taking the absolute value of the difference between the lateral-referenced ISOS and COST traces, i.e. |*ISOS*_*lateral*_ − *COST*_*lateral*_ |.

The local phase referencing approach relies on using a spatially localized visual stimulus (a circular disk; Fig. 2A&B, Protocol 2 in Methods) in an AO line-scan OCT system. This allows for changes in optical phase to be calculated for any given layer between the stimulated area and the surrounding non-stimulated area (see *Supplementary Section 1.2*). AO helped improve signal-to-noise ratio and hence phase stability for lateral differencing of these minute signals, while also localizing the responses to individual cones. Figure 2B displays representative ΔOPL maps for the four dominant outer retinal layers, averaged over 0.4-0.6 s post-stimulus, obtained by differencing the stimulated and non-stimulated areas. Within the stimulated regions, both near the fovea and at parafoveal eccentricity, the ISOS junction shows a vitread (towards the vitreous) displacement, COST shows a sclerad (towards the sclera) displacement, while the RPE exhibited negligible movement, within the noise floor of the measurement. The displacement of the ELM was eccentricity-dependent, with a notable vitread shift near the fovea but minimal change in the parafovea (Fig. 2B). The displacement of specific outer retinal layers near the fovea and in the parafovea is summarized in Fig. 2D with a schematic representation using a cartoon of the cone photoreceptor denoting the different compartments.

The time-varying ΔOPL traces for each layer, obtained by spatially averaging the ΔOPL signals across the stimulated regions, confirmed that these displacements initiated rapidly after stimulus onset and persisted throughout the measurement time window up to 2.5 s (Fig. 2C). Their kinetics were similar to the deformations of the encasing compartments seen in Fig 1. The displacements of the ELM, ISOS and COST were substantially larger near the fovea than in the parafovea, whereas the RPE remained measurably static in both regions. Notably, the traces contained small quasi-periodic oscillations (Fig. 2C), attributable to pulsatile motion of the underlying tissue that is essentially eliminated when these layer-specific displacements are subjected to axial referencing as in Fig. 1 and Fig. 2E. The bleach-dependence of these layer-specific displacements are provided in Supplementary Fig. S8.

It bears verifying whether the measurements undertaken with these two distinct phase referencing paradigms – axial and lateral – yield the consistent values. Given that the RPE exhibited negligible movement following visual stimulation (see pink curves in Fig. 2C), we first tested whether this layer could be treated as a stationary reference in the axial-referencing paradigm to obtain absolute displacements of the other outer retinal interfaces. In Fig. 2E, the axial-referenced separations: ISOS–RPE, COST–RPE, and ELM–RPE, closely resemble the corresponding unsigned absolute displacements of ISOS, COST, and ELM obtained from lateral referencing. Moreover, it suppresses the common-mode perturbations attributed to bulk tissue pulsatility that cancel out with axial differencing, yielding layer-pair traces that are minimally affected by the oscillatory modulation evident in the traces of Fig. 2C. Furthermore, we verified that the cone OS expansion computed from the difference between the lateral-referenced ISOS and COST traces (dashed blue curves in Fig. 2E), matched well with the cone OS expansion calculated using conventional axial-referencing (i.e. |*ISOS* − *COST*|; solid blue curves in Fig. 2E). In summary, Fig. 2 establishes the direction and magnitude of displacement for the dominant outer retinal interfaces – ELM, ISOS, COST and RPE and links them directly to the deformations of the respective compartments – inner segment (IS), outer segment (OS) and supracone space (SCS).

### Stimulus strength and eccentricity-dependence of outer retinal deformations

We next examined how the light-evoked displacements of the layers vary with stimulus strength and retinal eccentricity. Varying stimulus strength is expected to modulate the magnitude of the mechanical force generated within the cone OS during phototransduction, while retinal eccentricity reflects the natural variation in cone photoreceptor geometry, including OS length and IS diameter, and the surrounding retinal layer architecture^24,25^. Characterizing these dependencies provides insight into how intrinsic actuation and anatomical structure together shape the mechanical response of the outer retina.

Figure 2E showed that the RPE can serve as a stationary reference, enabling absolute displacements of the other interfaces to be recovered by axial referencing. Therefore, the dependence on stimulus strength and eccentricity was measured using the same axial referencing approach from data acquired across a 5-deg field of view with a coarse-scale ORG instrument described elsewhere^45^. This platform reduces the burden of maintaining fixation and enables higher-throughput, wide-field recordings within a single acquisition. Figure 3A shows a representative *en face* retinal image at the layer corresponding to the ISOS junction, spanning the fovea to 5° temporal eccentricity. Axial-referenced ΔOPL traces at foveal and parafoveal eccentricities (boxes in Fig. 3A) were extracted for five interface pairs – those referenced to the RPE (ELM–RPE, ISOS–RPE and COST– RPE, Figs. 3B-D), and pairs spanning the cone outer segment (ISOS – COST; Fig. 3E) and inner segment (ELM – ISOS; Fig. 3F). Three stimulus levels were tested in 4 subjects (see Protocol 1 in Methods). Overall, the magnitude of deformations exhibited a rapid post-stimulus rise (or fall) followed by a slower approach to a plateau, with the amplitude and rate of rise scaling with stimulus level (see Figs. 3B–F). As in Fig. 3C&3D, ΔOPL decreased in the supracone space (COST–RPE) but increased for the OS-SCS compartment taken as a whole (ISOS–RPE), consistent with the sclerad shift of COST and vitread shift of ISOS observed in Fig. 2. Larger amplitudes and faster rise times were consistently observed near the fovea across all compartments, consistent with the behavior seen for cone OS expansion here (Fig. 3E) and in prior reports^38^. Interestingly, the vitread displacement of the ELM (Fig. 2) and the corresponding increase in deformation of the ELM–RPE compartment (Fig. 3B) were evident near the fovea across all stimulus levels, but were largely absent in the parafovea.

**Figure 3.**
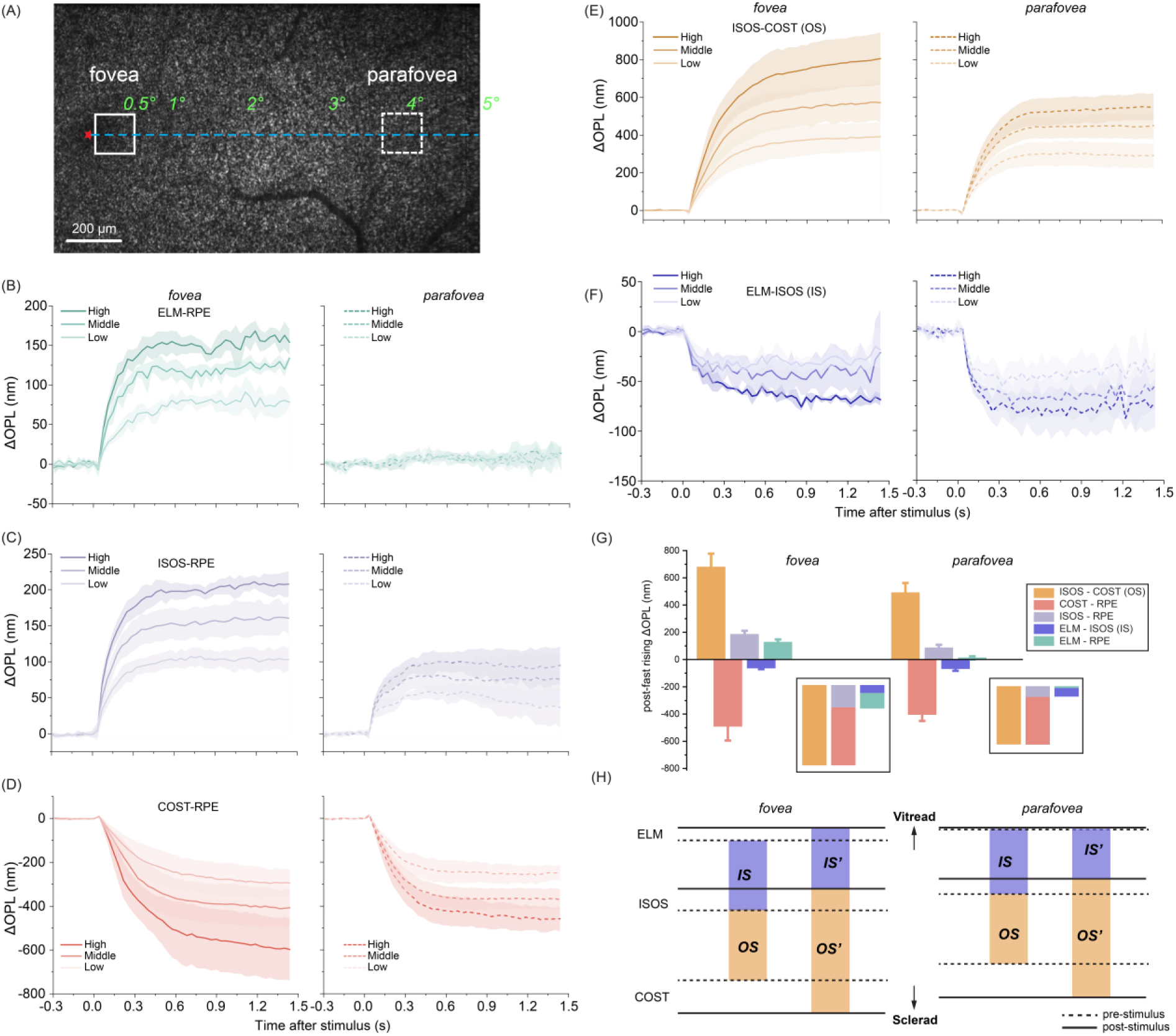
Stimulus strength and eccentricity dependence of light-evoked deformation of the outer retinal compartments. (A) En face structural image at the ISOS layer, obtained by segmenting a registered volume obtained from the CoORG instrument. Regions-of-interest used to compute the temporal evolution of the ΔOPL traces are outlined by white boxes. Fovea is marked by a red asterisk, and the temporal meridian is denoted by the blue dashed line. (B)-(F) The time evolution of deformation, expressed as ΔOPL vs. time, extracted from various retinal pairs for three stimulus bleach levels (high, middle, low, correspond to 14.1, 8.8, 4.4×10^6^ photons/µm^2^ on the retina, respectively) at the fovea (within 0.5°,left, solid traces) and parafovea (4°, right, dashed traces). Shaded envelopes indicate ±1 standard deviation (SD), obtained from measurements across 4 subjects. (G) ΔOPL amplitudes, obtained by averaging the time-varying ΔOPL traces over t = 0.4 – 0.6 s, are shown at the fovea (within 0.5°) & parafovea (4°). The black box insets show a comparison of the deformation magnitude between different partitions. Error bars denote ±1 SD, obtained from measurement across 10 subjects. (H) Schematic showing the light-evoked displacement of ELM, ISOS and COST and the deformation of the compartments they encase, i.e. the cone inner segment (IS) and outer segment (OS).

In a larger subject cohort (n = 10) at the highest stimulus level, the saturated amplitude of deformation (averaged between t = 0.4–0.6 s post-stimulus) is summarized across retinal eccentricities in Supplementary Fig. S9. The foveal and parafoveal results are shown in Fig 3G. The results reproduce the same sign and relative magnitude of layer displacements, with modest inter-subject variability. The insets (black boxes in Fig. 3G) show additivity across compartments: deformations measured over longer axial spans agree with the sum of their constituent segments (e.g., Δ(ISOS–RPE) ≈ Δ(ISOS–COST) + Δ(COST–RPE)), and this relationship holds across all layer combinations.

Whereas stimulus-evoked outer segment expansion and compression of the supracone space exhibited substantially larger amplitudes near the fovea than in the parafovea, the compression of the inner segment remained similar at both locations. The signed displacements of the COST and ISOS further reveal the relative contributions of these interfaces to the overall cone outer-segment elongation. Near the fovea, the vitread shift of the ISOS accounted for approximately 25–30% of the total COS elongation, whereas in the parafovea, this fraction decreased to ∼15–20%. The larger ISOS contribution near the fovea arises from both a compression *and* a vitread translation of the inner segment, which manifests as a measurable vitread displacement of the ELM. In contrast, the smaller contribution of ISOS in the parafovea is largely attributable to inner segment compression alone, as the ELM appears to be stationary at this eccentricity within the noise floor of the measurement. These patterns of compartment deformation and layer movements (Fig. 3H) indicate that the mechanical response of the outer retina varies systematically with retinal eccentricity and stimulus strength. A quantitative biomechanical interpretation of these deformations requires a framework that links the measured layer movement to variation in photoreceptor anatomy with eccentricity, the mechanical force generated within the outer segments by light stimulation, and the material properties of the surrounding retinal compartments.

### A hybrid analytical-finite element framework links retinal deformations to biomechanics

To link the measured deformations to the distinct material properties of the different retinal compartments and the eccentricity-dependent variation in photoreceptor anatomy, we developed a mechanical framework to assess how OS-driven deformation is distributed between the proximal ELM/inner-retina side and the distal SCS side. In the simplest representation, as shown in Fig. 4A, a cone photoreceptor was modeled as a series of axially-coupled springs (spring constants: *k*_IS_ and *k*_OS_) with proximal and distal supports (areal spring constants: *K*_A,IR_and *K*_A,SCS_, respectively). A light stimulus activates the elongation of the cone OS and provides the active drive for the spring series.

**Figure 4.**
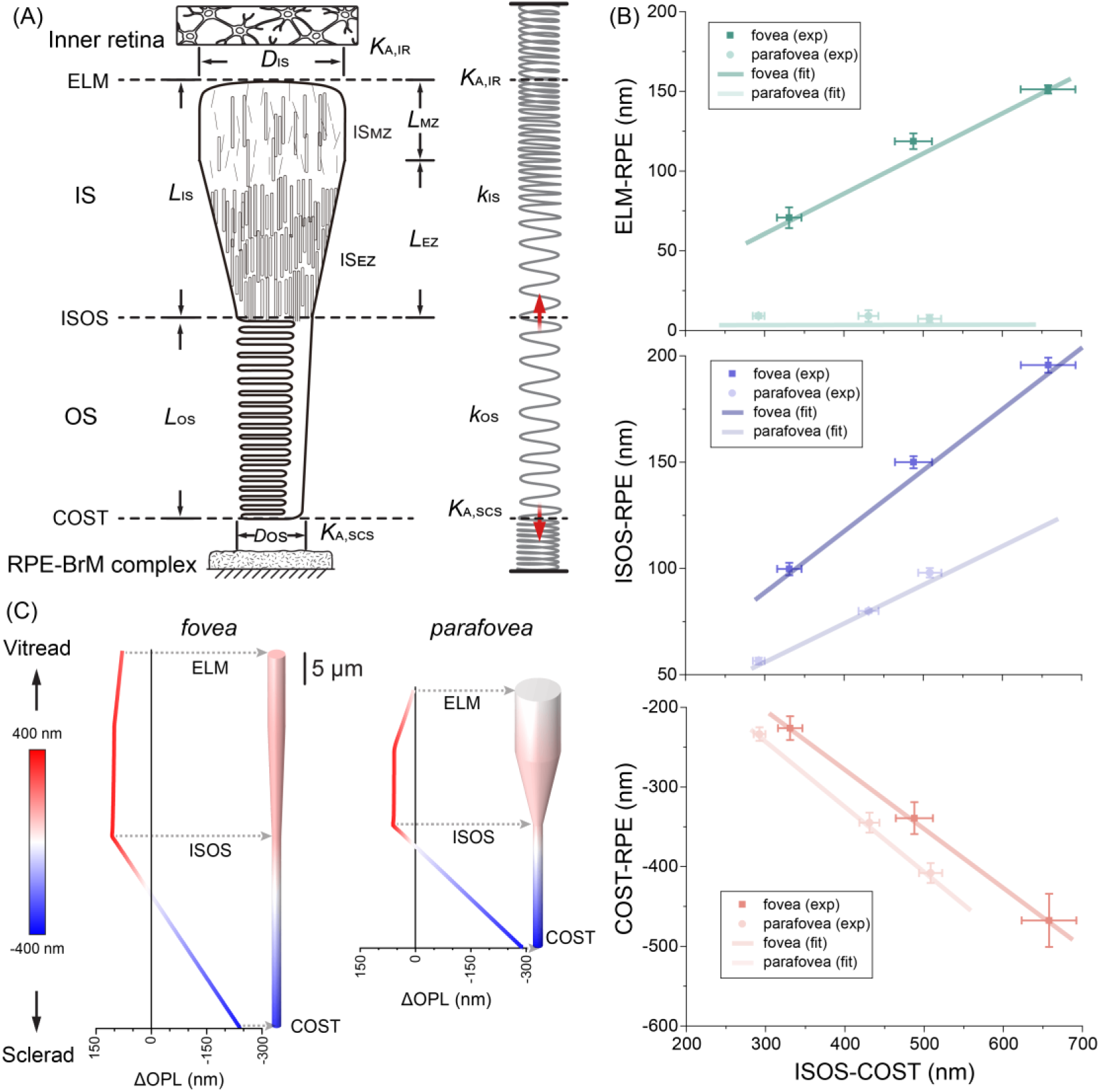
Comparison of the experimental measurements with a biomechanical model. (A) Schematic showing a single cone photoreceptor embedded between the inner retina and SCS. Each segment of the cone photoreceptor, including the myoid zone, ellipsoid zone and OS, is parameterized by its geometry and material properties (length *L*, cross-sectional area *A*, Young’s modulus *E*). The inner retina and SCS resist cone OS elongation by providing restoring force with areal spring constants of *K*_*A*,*IR*_ and *K*_*A,SCS*_, respectively. A linear helical spring-series is shown adjacent to the cone. Note that, for a helical spring, the spring constant scales inversely with the number of active coils. In the schematic, a segment with denser coils qualitatively represents a higher Young’s modulus. (B) At the fovea and parafovea, model predictions, quantified as ELM, ISOS, and COST displacements as a function of OS elongation (solid lines), matched the corresponding experimental measurements (ELM–RPE, ISOS–RPE, COST–RPE; solid circles) extracted from Fig. 3B-D. Error bars indicate ±1 standard deviation. obtained from measurements across 4 subjects. (C) COMSOL finite-element model predicted stimulus-evoked axial displacement field at the quasi-static, saturated state in foveal and parafoveal cones. The predicted deformation was converted to ΔOPL using a refractive index of 1.41^37^. Color bar shows magnitude and direction of displacement. Centerline profiles and pseudocolored renderings show a transition near the ISOS from smaller vitread motion (red) to larger sclerad motion toward COST (blue). A greater vitread displacement at ELM is evident in the foveal cone compared to the parafoveal cone.

To infer whether the observed deformation falls within a linear-elastic regime, we plotted the displacements of the ELM, ISOS, and COST (with respect to the RPE) against the elongation of the OS (ISOS-COST). Across the three stimulus strengths used in Fig. 3, the displacements varied approximately linearly with OS elongation (Fig. 4B, solid symbols). These observations are consistent with linear-elastic behavior and support the applicability of Hooke’s Law (*F = kx*), i.e. the linear dependence between force (*F*) and displacement (*x*), to the spring series under our experimental conditions. Accordingly, the OS elongation serves as a proxy for the light-induced driving force and scales linearly with the displacements of the ELM, ISOS and COST.

This linearity of the absolute displacements (output) with the stimulus (input) motivated the definition of two dimensionless fractions, *f*_ISOS_ and *f*_ELM_, which capture the fraction of OS elongation that is accounted for by the vitread ISOS motion, and the fraction of ISOS motion transmitted to ELM displacement, respectively. At each eccentricity, these fractions represent a drive-independent quantitative measure of the differential partitioning of deformations at the proximal and distal ends of the photoreceptor. The *f*_ISOS_ remained below 0.5 near the fovea and in the parafovea (mean ± standard deviation: 0.31 ± 0.05 and 0.19 ± 0.03, respectively), indicating that majority of the OS-driven deformation was directed distally towards the RPE. This distal bias was stronger in the parafovea, where a smaller fraction of OS elongation translated to a vitread ISOS motion, while majority of the elongation was accounted by the sclerad COST movement. In contrast, proximal partitioning of deformation differed sharply with eccentricity: *f*_ELM_ was high near the fovea (0.74 ± 0.04) but low in the parafovea (0.08 ± 0.04), indicating that near the fovea, a substantial fraction of proximal motion is transmitted to the ELM, whereas in the parafovea, the same motion is absorbed as a compression of the IS with little measurable ELM translation.

The measured values of *f*_ISOS_and *f*_ELM_, together with eccentricity dependent variation in cone geometry, were used to analytically solve the spring series model to retrieve the relative ratios of *K*_A,IR_, *K*_A,SCS_, and *k*_IS_. In addition, by anchoring the value *K*_A,IR_at the fovea, done independently using an inner-retina finite element (FE) model (see Methods, Supplementary Section 3.1), these relative stiffness ratios were placed on an absolute scale to retrieve the stiffnesses and elastic moduli of the different compartments. This treatment yielded an effective modulus of the SCS of 8.7 ± 2.0 Pa, consistent with its composition as hydrated extracellular space containing interphotoreceptor matrix and RPE microvilli, and is comparable in its stiffness to soft gel-like biological materials such as mucus ^46^. On the proximal end, the effective stiffness of the IS (*k*_IS_) was found to be almost twice as high in the fovea compared to the parafovea (*k*_IS,fov_ = (3.3 ± 0.7) × 10^−4^ N/m vs *k*_IS,para_ = (1.6 ± 0.3) × 10^−4^ N/m), suggested to arise from previously-noted differences in mitochondrial packing within the foveal and parafoveal cone IS^24,25^.

To estimate the relative stiffness contribution from the mitochondria-rich ellipsoid-zone (EZ) and the myoid-zone (MZ) of the IS, an analytical decomposition of the IS stiffness and a post hoc sensitivity analysis was performed (Methods and Supplementary Section 4.4). Inferred stiffnesses were converted to elastic moduli by considering the variation in cone geometry as a function of eccentricity (Fig. 4C). For a plausible range of EZ moduli, this analysis reveals the contrast in the MZ modulus between the fovea and parafovea (Supplementary Fig. S6). For example, using a representative ellipsoid zone modulus (*E*_*EZ*_= 40 kPa), consistent with reported estimates of mitochondrial stiffness ^47^, the inferred MZ modulus was *E*_*MZ*,0.25°_ = 0.67 ± 0.14 kPa near the fovea and *E*_*MZ*,4°_ = 0.05 ± 0.01 kPa in parafovea, i.e. a ∼14-fold difference between the two locations (see Supplementary Section 4.4).

Finally, using the inferred Young’s moduli of all the compartments from the analytical spring series model, together with measured eccentricity-dependent parameters of cone geometry, a forward COMSOL model was developed to simulate the biomechanical input-output response curve. The input was an initial stimulus-induced pressure in the OS, and the outputs were the deformations of the various compartments (Fig. 4C). Assuming a value (26 kPa) for the Young’s modulus of the OS ^48^, an input OS pressure in the range of 61-610 Pa was found to cause linear OS elongations of 100-1000 nm (Supplementary Fig. S7). Notably, a unique value of force generated in the OS was able to simultaneously reproduce the key experimental observations - near-stationary RPE, sclerad COST motion, vitread ISOS motion and eccentricity-dependent ELM movement – as well as the measured displacements in the three layers – ELM, ISOS and COST (Fig. 4B, solid lines).

The FE model extends the spring series model from layer-specific movements to enable the visualization of the entire deformation field across the compartments. As a representative example, Fig. 4C shows the deformations of a foveal and parafoveal cone photoreceptor for an input light-induced pressure at equilibrium of 213 Pa & 330 Pa, respectively, yielding an OS elongation of ∼350 nm, and a residual pressure towards the neighboring compartments of ∼10 Pa, implying that majority of the light-induced pressure is counterbalanced by the restoring force of OS itself. These values of OS pressure increase in the FE model that reproduce the measured compartmental deformations yield a quantitative estimate of the intrinsic mechanical actuation provided by light stimulation and provide important constraints for the underlying biophysical mechanisms generating the actuation (Supplementary Section 5).

### Altered light-evoked displacement and inter-layer coupling in retinitis pigmentosa

We next applied the framework to five patients with a clinical diagnosis of retinitis pigmentosa (RP), an inherited retinal dystrophy characterized by progressive photoreceptor degeneration^49^. In RP, a transition zone (TZ) demarcates an intact central region from peripheral degeneration and is defined by the loss of the COST reflection while ISOS and RPE signals remain detectable (Fig. 5A).

**Figure 5.**
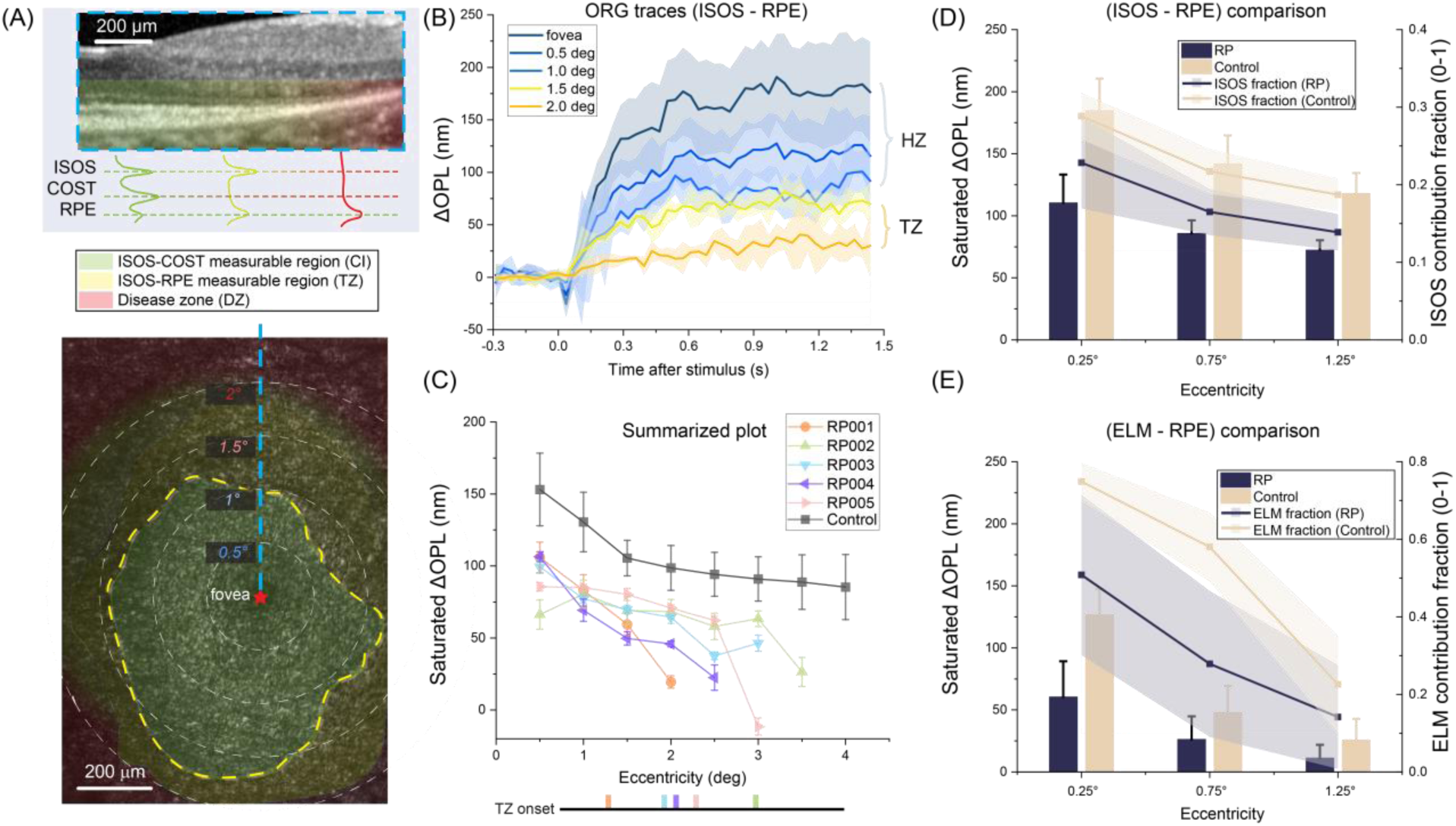
Clinical demonstration of ORG using COST-independent metrics and an ELM/ISOS coupling index in retinitis pigmentosa (RP). (A) Structural OCT/en face views delineating a preserved COST central region, a transition zone (TZ) with compromised COST but preserved ISOS and RPE, and an outer region where photoreceptor bands are not reliably measurable. CI: central island. DZ: disease zone. (B) Representative Δ(ISOS-RPE) at increasing eccentricities in response to visual stimuli with a photon density of 14.1 × 10^6^ photons⁄μm^2^.. Solid lines indicate mean responses; shaded regions indicate ±1 standard deviation (SD) range across measurements. (C) Summarized plot of ISOS-RPE amplitude (averaged time: t = 0.4 – 0.6s) as a function of eccentricity across 5 RP patients and 10 normal controls. (D) Summary of Δ(ISOS–RPE) amplitude (bars; mean ±SD) versus eccentricity for controls and RP, with overlaid ISOS contribution fraction (lines) plotted on the right axis (shaded area = ±SD). The ISOS contribution fraction was computed from saturated amplitudes as Δ(ISOS-RPE)/Δ(ISOS-COST) in regions where COST was measurable. (E) Summary of saturated Δ(ELM–RPE) amplitude (bars; mean ±SD) and ELM contribution fraction plotted on the right axis (lines; shaded area = ±SD) versus eccentricity for controls and RP. The ELM contribution fraction was defined as Δ(ELM-RPE)/ Δ(ISOS-RPE).

Light-evoked activity in the ISOS-RPE compartment remained measurable in both the central region and the TZ, including locations where the COST reflection was absent. The response amplitude decreased from the central retina to the TZ (Fig. 5B), consistent with the known spatial progression of disease. Compared to controls, vitread displacement of the ISOS was significantly reduced in all RP patients, even within the central retina where outer retinal structure appeared normal on clinical imaging (Fig. 5B&5C). The location of the TZ varied across the five subjects (Fig. 5C): in RP01, RP02 and RP03, two sampled locations fell within the TZ, whereas in RP04 and RP05, one location was within the TZ. At these locations inside the TZ, despite the loss of the COST reflection, vitread ISOS displacement remained detectable but was further reduced indicating residual but impaired light-evoked deformation.

We next assessed whether mechanical properties of the OS, IS and the inner retina were altered in regions that appeared structurally intact on OCT. To account for reduced light-evoked OS elongation in RP, ISOS displacement was expressed as a fraction of total OS elongation with an analogous normalization applied to ELM displacement, as defined earlier – *f*_*ISOS*_ and *f*_*ELM*_. Across subjects, both absolute ISOS displacement and *f*_*ISOS*_ was reduced in RP relative to controls (Fig. 5D). Similarly, ELM displacement was reduced in RP, with the most pronounced differences observed at the fovea (Fig. 5E).

Within the spring-series framework described above, the distribution of deformation across compartments is governed by their relative compliance, such that changes in the fractional metrics - *f*_*ISOS*_and *f*_*ELM*_ - reflect changes in effective stiffness and inter-layer coupling. A reduction in *f*_*ISOS*_is consistent with decreased compliance (i.e., increased effective stiffness) of the inner segment, while a reduction in *f*_*ELM*_ indicates altered mechanical coupling between the photoreceptor layer and the inner retina.

## Discussion

This article introduces a new paradigm for imaging retinal biomechanics in vivo. Common elastography techniques that estimate biomechanical properties from tissue motion often rely on externally-applied, controlled mechanical excitation (acoustic radiation force^50^, shaker^34^ etc.) with the most extensive applications in the eye to the anterior segment, particularly the cornea^27,51^. However, applications to the posterior segment remain technically challenging and mostly restricted to ex vivo preparations and animal models^50^. Brillouin microscopy offers a promising alternative for contact-free, optical assessment of viscoelastic properties, but similar to elastography, its application to posterior segment remains less developed than to the anterior segment ^30,28,29,31,52^. In the current approach, the key mechanical actuation central to elastography is provided by a light stimulus that triggers phototransduction which is also accompanied by a set of biophysical events that deform the photoreceptor and surrounding retinal structures. The spatially localized nature of the driving force originating in the OS and the distinct layer movements it produces across the thickness of the retina together allow linking the deformations to the specific biomechanical properties the retinal layers.

At its core, the current paradigm for imaging tissue biomechanics builds upon the emerging body of work in the field of ORG for measuring light-evoked outer retinal deformation^40–43,53,44,54,38,55,56^. While most studies have focused on the light-evoked deformation of the photoreceptor OS^38,40–42,45,55^, a few have reported deformations in surrounding compartments as well^44,57^. Although the underlying biophysical mechanism by which a light stimulus generates force and the resultant OS deformation remains incompletely understood, recent work in patients with bradyopsia places the underlying molecular mechanism of the cone OS elongation within the G-protein cascade at or prior to the GTP hydrolysis, catalyzed by the enzyme RGS9 (Regulator of G-protein Signaling 9)^58^. Importantly, the use of ORG for biomechanical inquiry does not require a complete understanding of the actuation mechanism and instead provides a framework that can be progressively refined as the molecular and biophysical basis of the deformation is clarified. It is worth noting, however, that this paradigm is currently restricted to a quasi-static, small deformation linear elastic regime, whereas retinal tissue may be expected to exhibit complex anisotropic, nonlinear, and viscoelastic behavior^10^. Thus, the FE model should be interpreted as estimating an elastic response at equilibrium, as defined by the light-evoked deformation plateau, while being insensitive to frequency dependent viscoelastic losses, damping, or dispersion that entail a measurement with dynamic or multifrequency elastographic modalities^59^. In this context, results from the FE model constrain the effective pressure at the plateau to lie in the range of ten to hundreds of Pa (Supplementary Fig. S7), for an OS elongation in the range of 100-1000nm, providing a quantitative estimate that informs both the mechanical modeling and the candidate molecular substrates of the underlying mechanism.

It has remained impossible thus far under the prevailing axial phase referencing paradigm used for ORG to attribute the deformation of a given compartment, say the OS, to the retinal interfaces encasing it. Given the heterogeneity in the material composition of the different retinal compartments, linking the deformations to biomechanical properties hinges on the knowledge of how each layer deforms in response to the driving force - a challenge overcome here by the introduction of local phase-referencing (Fig. 2A). Using spatially localized visual stimulation and differencing the phase emanating from stimulated and unstimulated areas, the absolute light-evoked movement was resolved in each of the major outer retinal layers (Fig. 2B&2C). By comparing these experimental results with a hybrid analytical spring-series and finite element model, the biomechanical properties of the different retinal compartments in the fovea and parafovea were deduced (Supplementary Table S4).

The majority of OS-driven deformation is directed sclerally (*f*_*ISOS*_ range: 0.31 ± 0.05 near the fovea and 0.19 ± 0.03 in the parafovea) consistent with the greater compliance estimated by the model for the distal sub-retinal side of the OS. In contrast, the proximal side was significantly stiffer and exhibited a marked difference between the fovea and parafovea. The difference is explained by distinct foveal and parafoveal cone IS diameter (2.8 µm vs. 6.5 µm), as well as the order-of-magnitude larger modulus of the myoid zone in foveal cones (Supplementary Fig. S6). The difference in IS diameters corresponds to an approximately 5-fold smaller cross-sectional area in foveal cones, which would result in greater axial stress for a given force and more efficient transmission of deformation toward the ELM. The difference in myoid zone modulus is qualitatively consistent with anatomical observations showing that mitochondria, known to exhibit high stiffness (modulus : ∼60 kPa^47^), extend proximally beyond the ellipsoid within the foveal cone IS^24,25^. Furthermore, the overlying inner retina in the parafovea appears to resist the propagation of motion, in contrast to the fovea where the inner retinal layers are displaced from the foveal pit.

Near the foveola, the efficient axial transfer of deformation is further supported by specialized microstructure. Cone axons in the Henle fiber layer are short and nearly aligned with the photoreceptor axis^60^, becoming more oblique in the parafovea before turning vertical again in the perifovea^60^. The Muller glial cells form a densely packed and vertically oriented structure termed the Muller cell cone, with at least one Muller cell per foveal cone^61,62^. Muller cells provide a continuous structural scaffold spanning the retina and contribute to mechanical coupling across layers. Their dense, columnar organization in the fovea along with the vertical Henle fibers may facilitate efficient axial transmission of deformation from the photoreceptor layer to the inner retina, with reduced lateral dissipation into surrounding tissue. In addition, the ILM is exceptionally thin in the fovea compared to the parafovea (138 nm vs. 3.7 µm)^63^, potentially reducing resistance to axial displacement in the fovea while constraining motion in the parafovea. Together, these anatomical specializations provide a structural basis for the efficient axial transfer of deformation in the fovea, and its attenuation in the parafovea.

Applying this framework to patients with RP revealed early functional and biomechanical alterations in disease. Light-evoked activity in the ISOS-RPE compartment remained measurable within the TZ (Fig. 5A), where the COST reflection is absent by definition, indicating that the deformation of this compartment can provide a measure of residual photoreceptor function that is independent of the reflective integrity of the COST (Fig. 5B-C), often known to be compromised even when the overlying photoreceptor retains intact inner segments and nuclei ^64^. Given that the transition zone represents the leading edge of degeneration^65–67^, a functional readout of these residual and vulnerable photoreceptors provides a sensitive biomarker for monitoring disease progression and therapeutic response in clinical trials^68,64,69^. Reduced fractional displacements (*f*_*ISOS*_ and *f*_*ELM*_,Fig. 5D-E), particularly at the fovea, suggest altered IS compliance and weakened mechanical coupling between the photoreceptors and the inner retina, even in regions that appear structurally intact on clinical imaging. These mechanical changes may reflect underlying alterations in cellular and tissue composition, including mitochondrial remodeling within the IS associated with metabolic stress^70^, as well as gliosis-associated changes in the inner retina that may modify mechanical coupling between photoreceptors and surrounding tissue^15,71^. Together, these findings provide promising evidence for functional impairment and biomechanical remodeling at early stages of disease. More broadly, this framework may be applicable to a range of retinal disorders where structural integrity and tissue biomechanics are altered^15,72,73^, including AMD^74^, vitreoretinal interface disorders^16^, and glaucoma^17^. By leveraging intrinsic optical actuation to probe both function and biomechanical properties in vivo, this approach introduces a new class of non-invasive elastography in the living human retina with direct relevance for clinical assessment of retinal disease.

## Methods

### Study participants

Study volunteers (n=10, ages 19–42 years old (yo), mean: 29 yo) with no known ocular conditions were recruited. Five patients (ages 23–60 years old (yo), mean: 35 yo) with a diagnosis of retinitis pigmentosa (RP) based on clinical presentation and family history were also recruited from the Roger and Angie Karalis Johnson Retina Center, Department of Ophthalmology at University of Washington. The characteristics of RP subjects are shown in Table S5. Research protocols were approved by an Institutional Review Board at the University of Washington (IRB#STUDY00002923). The study adhered to the tenets of the Declaration of Helsinki and written informed consent was obtained from all participants prior to study participation.

### System setup

Two custom-built line-scan spectral-domain OCT platforms were used to acquire retinal responses to light stimuli: an AO-OCT system^75^ for cellular-resolution measurements and for cone spectral classification and a coarse-scale ORG system without an AO module (CoORG)^45^ for larger field-of-view recordings. At their core, both instruments are based on a free-space, line-field configuration, where a cylindrical lens creates a linear imaging field on the retina and the backscattered light is captured in a high-speed 2D CMOS camera after diffraction from 1-D grating, such that in a single camera snapshot, an entire 2D-cross-sectional retinal image (B-scan) is acquired. The linear imaging field is scanned on the retina by a 1-D galvanometer scanner for volumetric OCT acquisition. The two instruments – CoORG and AO-OCT – are described previously. Key parameters include the imaging center wavelength and bandwidth - 820 ± 40 nm and 840 ± 25 nm, respectively for AO-OCT and CoORG. The imaging field of view with AO-OCT was adjustable up to 2° × 2°, while for CoORG the maximum field was 5° × 3°. The specific fields-of-view used in the experiments are given in the next section. A separate channel setup in Maxwellian view provided the stimulus for ORG and its spectral characteristics could be selected as per the experimental protocol, detailed below. To produce localized stimulation, a pinhole was inserted into the AO-OCT stimulus channel (Fig. 6A) and its spatial extent was measured using a model eye consisting of a lens and a reticle target placed at its focus as the artificial retina. The resulting localized and full-field stimulation patterns are shown in Figs. 6B and 6C.

**Figure 6.**
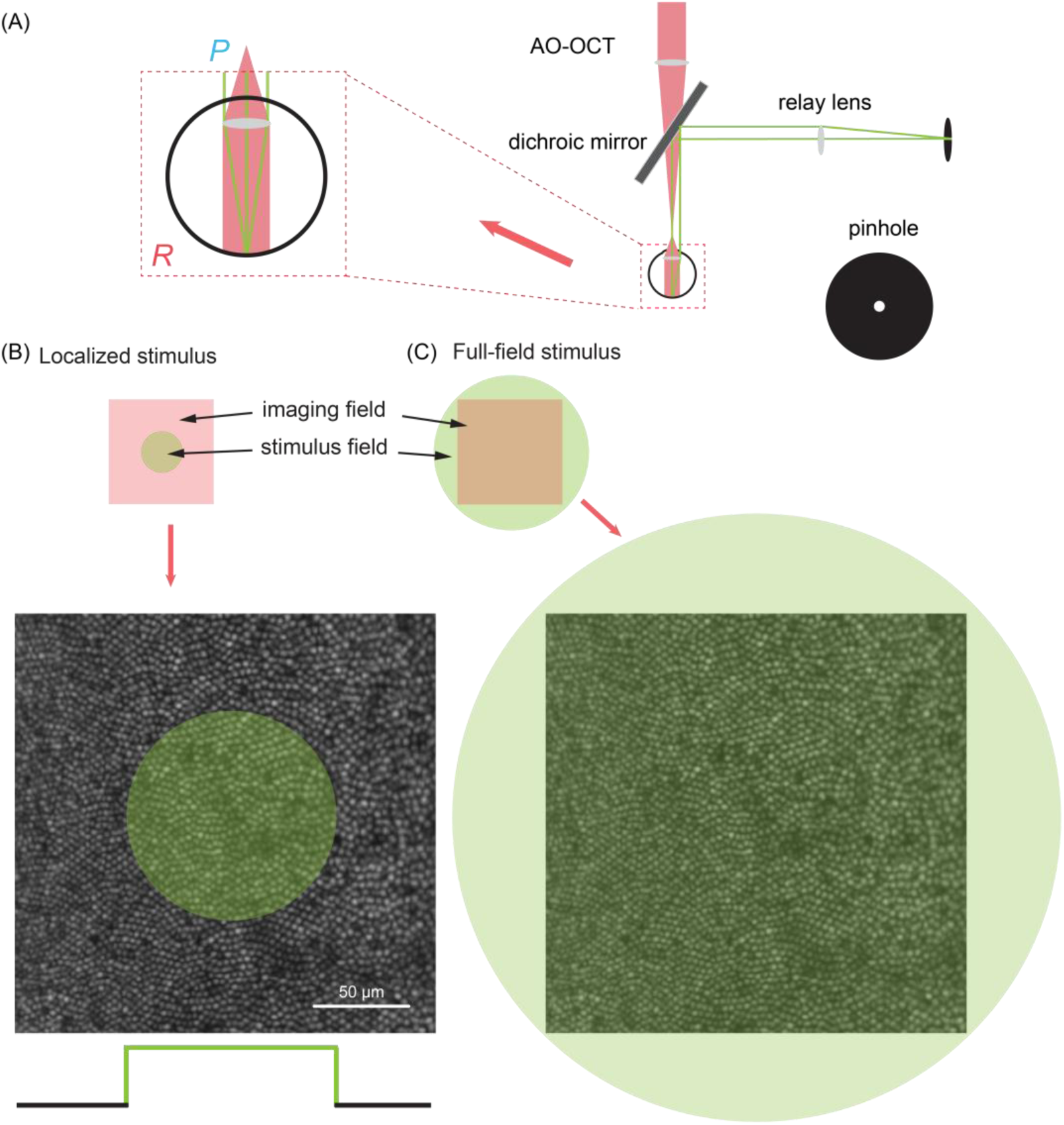
Stimulus delivery configurations for localized lateral versus full-field axial referencing. (A) Optical layout used to deliver the visual stimulus during AO-OCT acquisition. An LED source is shaped by a 600 µm diameter pinhole placed at a retinal conjugate plane in localized stimulus mode to control the lateral extent of the stimulus at the retina. P: Pupil plane; R: Retinal plane. The stimulus beam is combined with the OCT scanning beam via a dichroic mirror and co-aligned at the retina. The inset (“zoom-in”) illustrates how the pinhole defines a confined illumination spot, enabling a stimulated region and an adjacent non-stimulated reference region within the same field of view. (B) Localized stimulus configuration used for local referencing: in the cartoon schematic, the AO-OCT imaging field is shown as the pink square, while the small circular green region indicates the stimulus field. The stimulus is restricted to a localized retinal region within the imaging filed, while the surrounding area remains unilluminated. The unilluminated area serves as a control for removing common-mode phase drifts and bulk motion by differencing stimulated versus non-stimulated regions in the 3D OCT volume. (C) Full-field stimulus configuration used for axial referencing: the stimulus illuminates the full imaging field (green overlay both in cartoon and *en face* image).

### Optoretinography experiment protocol

For the healthy cohort, imaging was performed in either of the two eyes. Before the ORG experiments, pupils were pharmacologically dilated and cyclopleged using 1% tropicamide (Akorn Inc.). Participants underwent 1 min of dark adaptation prior to each recording. Each ORG acquisition consisted of 50 sequential OCT volumes, with a visual stimulus delivered after 10 volumetric scans. Between 5-10 acquisitions, each consisting of 50 volumes were acquired. Three ORG recording protocols –implemented using either the CoORG system or the AO-OCT system with different visual stimulation configurations, fields of view and speed – are detailed below. Clinical participants were imaged in the right eye using the same general procedures as the healthy cohort. To improve data reliability given reduced fixation stability, the number of repeat acquisitions was increased to 8–12 per subject. To limit total imaging time and reduce fatigue-related fixation instability, patient imaging was restricted to the brightest stimulus from *protocol 1*.

In *protocol 1* using CoORG system, OCT volumes were recorded over a 5° × 3° field of view at a B-scan rate of 16 kHz, corresponding to a volume rate of 32 Hz for 500 B-scans. Three nominal stimulus strengths were used – 4.4, 8.8, 14.1 × 10^6^ photons⁄μm^2^ on the retina– at a stimulus wavelength of 532 ± 5 nm. Photon density at the retina was computed by incorporating the measured stimulus spectral power distribution at the cornea, published values of anterior segment absorption and retinal image scaling, assuming a 7 mm pupil diameter and 24 mm axial length^45^.

*Protocols 2 & 3* were performed using the AO-OCT system, in which volumes were recorded over a 0.95° × 1.2° field of view at a B-scan rate of 12 kHz with 600 B-scans per volume yielding a volume rate of 20 Hz. *Protocol 2* adopted a localized stimulus (wavelength: 532 ± 5 nm, photon density on the retina: 2.2, 8.8, × 10^6^ photons⁄μm^2^) achieved by adding a pinhole into the stimulus path (see Fig. 6). *Protocol 3* used a full-field stimulus, centered at 660 ± 5 nm (18.8× 10^6^ photons⁄μm^2^). The latter wavelength for identifying cone spectral classes following a method described previously ^76^.

### Image processing and phase trace extraction

Raw spectral interferograms were reconstructed into complex-valued OCT volumes using standard spectral-domain processing (spectral calibration, k-space linearity remapping, and Fourier transform). Volumes affected by blinks or poor fixation were excluded. Outer-retinal interfaces were segmented in each dataset based on a previous publication^77^. For consistent visualization across eccentricities and to reduce curvature-induced confounds, volumes were flattened to the ISOS interface prior to extracting *en face* maps and extracting regions of interest centered at either the fovea or the parafovea. The repeated acquisitions were co-registered to compensate for eye movement based on the enface ISOS layer image using a strip-based image registration algorithm^78^. Combining the axial (z) shifts obtained from the layer segmentations with the 2D (xy) shifts from the strip-based algorithm, a complex-valued OCT volume series over time is obtained that is spatially registered in all three dimensions.

To achieve reliable and sensitive phase measurements in OCT, a common practice involves selecting a reference region within the sample and extracting phase dynamics relative to this reference. In this study, we employed two methods for extracting phase dynamics: the first method calculates the phase difference between layer pairs along the same A-line, denoted as axial-referencing (see Supplementary Section 1.1); the second method differences phase changes from regions within and outside the stimulated area, denoted as lateral-referencing (see Supplementary Section 1.2). The axial referencing method is used in all current implementations of ORG using phase-resolved OCT and captures the *relative* phase change between the layer pair along the A-line evoked by the stimulus. The lateral-referencing method assumes that the region illuminated by the stimulus undergoes a light-evoked phase-change in addition to unwanted phase drifts, while the unilluminated region is subject to the same phase drifts but does not undergo a light-evoked change. Subtracting the common mode noise attributed to the phase drifts between the two regions yields a measure of the stimulus-evoked response, providing the *absolute* magnitude of light-evoked response across the retinal thickness. The extracted phase signals were finally unwrapped along the temporal dimension and converted to changes in optical path length (ΔOPL) based on the central wavelength of the OCT system.

### Analytical and finite element modeling of retinal deformations to retrieve biomechanics

The stimulus-evoked deformation of the cone photoreceptor and its immediate surroundings were modeled as a quasi-static, 1D axial mechanical spring series. The active spring element was the cone OS, whose elongation Δ(ISOS–COST) provided the active drive. The COST boundary was resisted by a SCS areal spring constant *K*_A,SCS_, whereas the ELM boundary was resisted by an inner-retinal areal spring constant *K*_A,IR_. The cone IS was first treated as having a single effective axial stiffness *k*_IS_; decomposition into myoid-zone (MZ) and ellipsoid-zone (EZ) contributions were performed only after *k*_IS_ had been inferred.

The measured layer displacements were reduced to two dimensionless fractions, 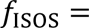 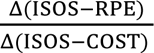 and 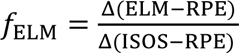 . The fraction, *f*_ISOS_, reports proximal-versus-distal partitioning of OS elongation, i.e. the magnitude of ISOS movement that accounts for the entire OS elongation. The fraction, *f*_*ELM*_, reports the extent to which the proximal motion of the ELM is accounted for by IS movement or compression. These fractions were estimated from the corresponding displacement pairs across the three stimulus levels, so that inferences on deformation distribution could be made independent of drive amplitude.

Closed-form expressions relating *f*_ISOS_and *f*_ELM_ to *K*_A,IR_, *K*_A,SCS_, *k*_IS_, and eccentricity-specific cone geometry are given in the Supplementary Section 2.3. These equations provide relative stiffness relationships obtained from the layer displacements. To place these relative stiffness values on an absolute scale, we constructed an independent inner-retina finite-element model in COMSOL Multiphysics v6.3 (Structural Mechanics Module), constrained by measured inner-retinal deformation, retinal layer thicknesses, and published Young’s moduli, to estimate the inner retinal stiffness, *K*_A,IR_, near the fovea (Supplementary Section 3.1). This anchor was then inserted into the analytical solution to obtain absolute values of *K*_A,SCS_ and *k*_IS_ (Supplementary Section 4.2-4.3) at the fovea. Absolute values of *K*_A,SCS_ at the parafovea were obtained under the working assumption that effective SCS support did not vary substantially between the sampled eccentricities.

This hybrid analytical-FE framework considers the retinal constituents either at the granular single cell level or at a gross tissue level. The spring series analytical model was formulated at the scale of a single cone, consistent with the observation that light-evoked deformations in the cone IS, OS and SCS can be resolved at the level of individual photoreceptors, enabling segregation of responses across spectral types - long, middle and short-wavelength cones (Supplementary Fig. S2C). On the other hand, deformation of the proximal compartment, ELM–RPE, does not exhibit cellular-scale specificity (Supplementary Fig. S2E). This arises because the cone spectral type-specific responses, which were separable at the OS, IS and SCS, become indistinguishable at the ELM boundary. This loss of cone-type specificity is interpreted to reflect the deformation of the shared ELM and inner-retinal scaffold (see Supplementary Section 3.1) rather than individual cones. Accordingly, the boundary condition proximal to the ELM was modeled at a tissue level.

Once the composite *k*_IS_was obtained, the IS was decomposed into MZ and EZ contributions by sweeping plausible *E*_EZ_values and solving for the corresponding *E*_MZ_using the cone geometry (Supplementary Section 4.4, Table S3). This sensitivity analysis was used to illustrate plausible and relative stiffness contributions by the two compartments. A separate COMSOL calibration related *K*_A,SCS_ to an effective SCS modulus, *E*_SCS_ (Supplementary Section 3.2).

Finally, a full COMSOL model was populated with the final parameter set and used for forward validation and visualization of multilayer deformation (Fig. 4B, Supplementary Section 3.3). The IS and OS are simply built by a tapering frustum and a cylinder with corresponding geometrical parameters derived from literature. Estimation of the total light-induced OS pressure at equilibrium was performed separately using OS stiffness from literature^48^; only the residual transmitted pressure entered the analytical spring series model (see Supplementary Section 5). Eccentricity-specific cone geometry was incorporated explicitly into the full COMSOL model because axial stiffness scales with geometry; accordingly, near foveal and perifoveal cones were modeled with different IS diameters and OS/IS (MZ+EZ) lengths (Table S3) based on anatomical studies and high-resolution imaging^25,79^. The summary of inferred biomaterial parameters such as Young’s modulus, stiffness of different outer retinal compartments, as well as areal spring constants of boundaries are reported in Supplementary Table S4.

## Acknowledgements

1. T. Liu acknowledges the support of the SPIE Franz-Hillenkamp Postdoctoral Fellowship. D. Mustafi acknowledges support from the Foundation Fighting Blindness. T. Ling acknowledges support of the National Research Foundation, Singapore for the NRF Fellowship Award (NRF-NRFF14-2022-0005), the Startup Grant from Nanyang Technological University, the seed funding programme under NMRC Centre Grant – Singapore Imaging Eye Network (SIENA) (NMRC/CG/C010A/2017), and the National Medical Research Council (MOH-001748-00). D. Mustafi acknowledges support from the Foundation Fighting Blindness. R. Sabesan acknowledges support from National Institutes of Health grants U01EY032055, EY029710, P30EY001730, unrestricted grant from the Research to Prevent Blindness; George and Martina Kren Professorship in Vision Research, Dawn’s Light Foundation, Kren Engineering in Medicine Initiative.

## References

1. Humphrey, J. D., Dufresne, E. R. & Schwartz, M. A. Mechanotransduction and extracellular matrix homeostasis. Nat. Rev. Mol. Cell Biol. 15, 802–812 (2014).

2. DuFort, C. C., Paszek, M. J. & Weaver, V. M. Balancing forces: architectural control of mechanotransduction. Nat. Rev. Mol. Cell Biol. 12, 308–319 (2011).

3. Guillot, C. & Lecuit, T. Mechanics of Epithelial Tissue Homeostasis and Morphogenesis. Science 340, 1185–1189 (2013).

4. Hannezo, E. & Heisenberg, C.-P. Mechanochemical feedback loops in development and disease. Cell 178, 12–25 (2019).

5. Phillip, J. M., Aifuwa, I., Walston, J. & Wirtz, D. The mechanobiology of aging. Annu. Rev. Biomed. Eng. 17, 113–141 (2015).

6. Handorf, A. M., Zhou, Y., Halanski, M. A. & Li, W.-J. Tissue stiffness dictates development, homeostasis, and disease progression. Organogenesis 11, 1–15 (2015).

7. Guertler, C. A. et al. Mechanical properties of porcine brain tissue in vivo and ex vivo estimated by MR elastography. J. Biomech. 69, 10–18 (2018).

8. Doyley, M. M. & Parker, K. J. Elastography: general principles and clincial applications. Ultrasound Clin. 9, 1 (2014).

9. Chatelin, S., Constantinesco, A. & Willinger, R. Fifty years of brain tissue mechanical testing: from in vitro to in vivo investigations. Biorheology 47, 255–276 (2010).

10. Ferrara, M. et al. Biomechanical properties of retina and choroid: a comprehensive review of techniques and translational relevance. Eye 35, 1818–1832 (2021).

11. Serai, S. D. & Yin, M. MR elastography of the abdomen: basic concepts. In Preclinical MRI of the Kidney: Methods and Protocols 301–323 (Springer, 2021).

12. Weaver, J. B. et al. Brain mechanical property measurement using MRE with intrinsic activation. Phys. Med. Biol. 57, 7275–7287 (2012).

13. Chung, S., Breton, E., Mannelli, L. & Axel, L. Liver stiffness assessment by tagged MRI of cardiac-induced liver motion. Magn. Reson. Med. 65, 949–955 (2011).

14. Harouni, A. A. et al. Assessment of liver fibrosis using fast strain-encoded MRI driven by inherent cardiac motion. Magn. Reson. Med. 74, 106–114 (2015).

15. Jones, B. W. et al. Retinal remodeling in human retinitis pigmentosa. Exp. Eye Res. 150, 149–165 (2016).

16. Davis, J. T., Wen, Q., Janmey, P. A., Otteson, D. C. & Foster, W. J. Müller cell expression of genes implicated in proliferative vitreoretinopathy is influenced by substrate elastic modulus. Invest. Ophthalmol. Vis. Sci. 53, 3014–3019 (2012).

17. Bellezza, A. J. et al. Deformation of the lamina cribrosa and anterior scleral canal wall in early experimental glaucoma. Invest. Ophthalmol. Vis. Sci. 44, 623–637 (2003).

18. Ugarte, M., Hussain, A. & Marshall, J. An experimental study of the elastic properties of the human Bruch’s membrane-choroid complex: relevance to ageing. Br. J. Ophthalmol. 90, 621–626 (2006).

19. L’Abbate, D., Prescott, K., Geraghty, B., Kearns, V. R. & Steel, D. H. Biomechanical considerations for optimising subretinal injections. Surv. Ophthalmol. 69, 722–732 (2024).

20. Irigoyen, C. et al. Subretinal injection techniques for retinal disease: a review. J. Clin. Med. 11, 4717 (2022).

21. Butterwick, A. et al. Effect of shape and coating of a subretinal prosthesis on its integration with the retina. Exp. Eye Res. 88, 22–29 (2009).

22. Holz, F. G. et al. Subretinal photovoltaic implant to restore vision in geographic atrophy due to AMD. N. Engl. J. Med. 394, 232–242 (2026).

23. Curcio, C. A., Sloan, K. R., Kalina, R. E. & Hendrickson, A. E. Human photoreceptor topography. J. Comp. Neurol. 292, 497–523 (1990).

24. Hoang, Q. V., Linsenmeier, R. A., Chung, C. K. & Curcio, C. A. Photoreceptor inner segments in monkey and human retina: Mitochondrial density, optics, and regional variation. Vis. Neurosci. 19, 395–407 (2002).

25. Spaide, R. F. & Curcio, C. A. ANATOMICAL CORRELATES TO THE BANDS SEEN IN THE OUTER RETINA BY OPTICAL COHERENCE TOMOGRAPHY: Literature Review and Model. Retina 31, 1609–1619 (2011).

26. Burns, S. A., Elsner, A. E., Sapoznik, K. A., Warner, R. L. & Gast, T. J. Adaptive optics imaging of the human retina. Prog. Retin. Eye Res. 68, 1–30 (2019).

27. Ramier, A. et al. In vivo measurement of shear modulus of the human cornea using optical coherence elastography. Sci. Rep. 10, 17366 (2020).

28. Scarcelli, G., Besner, S., Pineda, R., Kalout, P. & Yun, S. H. In vivo biomechanical mapping of normal and keratoconus corneas. JAMA Ophthalmol. 133, 480–482 (2015).

29. Besner, S., Scarcelli, G., Pineda, R. & Yun, S.-H. In vivo Brillouin analysis of the aging crystalline lens. Invest. Ophthalmol. Vis. Sci. 57, 5093–5100 (2016).

30. Scarcelli, G. & Yun, S. H. In vivo Brillouin optical microscopy of the human eye. Opt. Express 20, 9197 (2012).

31. Ambekar, Y. S. et al. Characterization of retinal biomechanical properties using Brillouin microscopy. J. Biomed. Opt. 25, (2020).

32. Larin, K. V. & Sampson, D. D. Optical coherence elastography – OCT at work in tissue biomechanics [Invited]. Biomed. Opt. Express 8, 1172 (2017).

33. He, Y., et al. Confocal Shear Wave Acoustic Radiation Force Optical Coherence Elastography for Imaging and Quantification of the *In Vivo* Posterior Eye. IEEE J. Sel. Top. Quantum Electron. 25, 1–7 (2019).

34. Qian, X., et al. *In vivo* evaluation of posterior eye elasticity using shaker-based optical coherence elastography. Exp. Biol. Med. 245, 282–288 (2020).

35. Zevallos-Delgado, C. et al. Acoustic radiation force optical coherence elastography of the crystalline lens: safety. Transl. Vis. Sci. Technol. 13, 36–36 (2024).

36. Arshavsky, V. Y., Lamb, T. D. & Pugh, E. N. G Proteins and Phototransduction. Annu. Rev. Physiol. 64, 153–187 (2002).

37. Pandiyan, V. P., Nguyen, P. T., Pugh, E. N. & Sabesan, R. Human cone elongation responses can be explained by photoactivated cone opsin and membrane swelling and osmotic response to phosphate produced by RGS9-catalyzed GTPase. Proc. Natl. Acad. Sci. 119, e2202485119 (2022).

38. Pandiyan, V. P. et al. The optoretinogram reveals the primary steps of phototransduction in the living human eye. Sci. Adv. 6, eabc1124 (2020).

39. Choma, M. A., Ellerbee, A. K., Yang, C., Creazzo, T. L. & Izatt, J. A. Spectral-domain phase microscopy. Opt. Lett. 30, 1162–1164 (2005).

40. Hillmann, D. et al. In vivo optical imaging of physiological responses to photostimulation in human photoreceptors. Proc. Natl. Acad. Sci. 113, 13138–13143 (2016).

41. Zhang, F., Kurokawa, K., Lassoued, A., Crowell, J. A. & Miller, D. T. Cone photoreceptor classification in the living human eye from photostimulation-induced phase dynamics. Proc. Natl. Acad. Sci. 116, 7951–7956 (2019).

42. Azimipour, M. et al. Optoretinogram: optical measurement of human cone and rod photoreceptor responses to light. Opt. Lett. 45, 4658–4661 (2020).

43. Tomczewski, S. et al. Light-adapted flicker optoretinograms captured with a spatio-temporal optical coherence-tomography (STOC-T) system. Biomed. Opt. Express 13, 2186–2201 (2022).

44. Li, H. et al. Optoretinography reveals rapid rod photoreceptor movement upon rhodopsin activation. Light Sci. Appl. 15, 58 (2026).

45. Jiang, X., Liu, T., Pandiyan, V. P., Slezak, E. & Sabesan, R. Coarse-scale optoretinography (CoORG) with extended field-of-view for normative characterization. Biomed. Opt. Express 13, 5989 (2022).

46. Guimarães, C. F., Gasperini, L., Marques, A. P. & Reis, R. L. The stiffness of living tissues and its implications for tissue engineering. Nat. Rev. Mater. 5, 351–370 (2020).

47. Li, Y., Honda, S., Iwami, K., Ohta, Y. & Umeda, N. Analysis of mitochondrial mechanical dynamics using a confocal fluorescence microscope with a bent optical fibre. J. Microsc. 260, 140–151 (2015).

48. Qu, Y. et al. Quantified elasticity mapping of retinal layers using synchronized acoustic radiation force optical coherence elastography. Biomed. Opt. Express 9, 4054 (2018).

49. Hartong, D. T., Berson, E. L. & Dryja, T. P. Retinitis pigmentosa. The Lancet 368, 1795–1809 (2006).

50. Qu, Y. et al. In Vivo Elasticity Mapping of Posterior Ocular Layers Using Acoustic Radiation Force Optical Coherence Elastography. Investig. Opthalmology Vis. Sci. 59, 455 (2018).

51. Lan, G., Aglyamov, S. R., Larin, K. V. & Twa, M. D. In vivo human corneal shear-wave optical coherence elastography. Optom. Vis. Sci. 98, 58–63 (2021).

52. Kabakova, I. et al. Brillouin microscopy. Nat. Rev. Methods Primer 4, 8 (2024).

53. Wongchaisuwat, N. et al. Optical coherence tomography split-spectrum amplitude-decorrelation optoretinography detects early central cone photoreceptor dysfunction in retinal dystrophies. Transl. Vis. Sci. Technol. 13, 5–5 (2024).

54. Zhang, P. et al. In vivo optophysiology reveals that G-protein activation triggers osmotic swelling and increased light scattering of rod photoreceptors. Proc. Natl. Acad. Sci. 114, (2017).

55. Ma, G., Son, T., Kim, T.-H. & Yao, X. Functional optoretinography: concurrent OCT monitoring of intrinsic signal amplitude and phase dynamics in human photoreceptors. Biomed. Opt. Express 12, 2661–2669 (2021).

56. Cooper, R. F., Tuten, W. S., Dubra, A., Brainard, D. H. & Morgan, J. I. Non-invasive assessment of human cone photoreceptor function. Biomed. Opt. Express 8, 5098–5112 (2017).

57. Tan, B. et al. Light-evoked deformations in rod photoreceptors, pigment epithelium and subretinal space revealed by prolonged and multilayered optoretinography. Nat. Commun. 15, 5156 (2024).

58. Weiss, C. E. et al. Optoretinography in R9AP-bradyopsia reveals the essential role of G-protein signaling in the human cone elongation response.

59. Parker, K. J., Szabo, T. & Holm, S. Towards a consensus on rheological models for elastography in soft tissues. Phys. Med. Biol. 64, 215012 (2019).

60. Ramtohul, P., Cabral, D., Sadda, S., Freund, K. B. & Sarraf, D. The OCT angular sign of Henle fiber layer (HFL) hyperreflectivity (ASHH) and the pathoanatomy of the HFL in macular disease. Prog. Retin. Eye Res. 95, 101135 (2023).

61. Gass, J. D. M. Müller Cell Cone, an Overlooked Part of the Anatomy of the Fovea Centralis: Hypotheses Concerning Its Role in the Pathogenesis of Macular Hole and Foveomacular Retinoschisis. Arch. Ophthalmol. 117, 821 (1999).

62. Masri, R. A., Greferath, U., Fletcher, E. L., Martin, P. R. & Grünert, U. Immunohistochemistry and Spatial Density of Müller Cells in the Human Fovea. Invest. Ophthalmol. Vis. Sci. 66, 46 (2025).

63. Henrich, P. B. et al. Nanoscale Topographic and Biomechanical Studies of the Human Internal Limiting Membrane. Investig. Opthalmology Vis. Sci. 53, 2561 (2012).

64. Wendel, B. J. et al. Multimodal High-Resolution Imaging in Retinitis Pigmentosa: A Comparison Between Optoretinography, Cone Density, and Visual Sensitivity. Invest. Ophthalmol. Vis. Sci. 65, 45 (2024).

65. Hood, D. C., Lazow, M. A., Locke, K. G., Greenstein, V. C. & Birch, D. G. The transition zone between healthy and diseased retina in patients with retinitis pigmentosa. Invest. Ophthalmol. Vis. Sci. 52, 101–108 (2011).

66. Aleman, T. S. et al. Retinal Laminar Architecture in Human Retinitis Pigmentosa Caused by *Rhodopsin* Gene Mutations. Investig. Opthalmology Vis. Sci. 49, 1580 (2008).

67. Sun, L. W. et al. Assessing photoreceptor structure in retinitis pigmentosa and Usher syndrome. Invest. Ophthalmol. Vis. Sci. 57, 2428–2442 (2016).

68. Lassoued, A., et al. Cone photoreceptor dysfunction in retinitis pigmentosa revealed by optoretinography. Proc. Natl. Acad. Sci. 118, e2107444118 (2021).

69. Liu, T. et al. Longitudinal Changes in Optoretinography Provide an Early and Sensitive Biomarker of Outer Retinal Disease. Am. J. Ophthalmol. 277, 375–386 (2025).

70. Jiang, K. et al. Multiomics analyses reveal early metabolic imbalance and mitochondrial stress in neonatal photoreceptors leading to cell death in *Pde6brd1/rd1* mouse model of retinal degeneration. Hum. Mol. Genet. 31, 2137–2154 (2022).

71. Hippert, C. et al. Müller Glia Activation in Response to Inherited Retinal Degeneration Is Highly Varied and Disease-Specific. PLOS ONE 10, e0120415 (2015).

72. Marc, R. E., Jones, B. W., Watt, C. B. & Strettoi, E. Neural remodeling in retinal degeneration. Prog. Retin. Eye Res. 22, 607–655 (2003).

73. Cuenca, N. et al. Cellular responses following retinal injuries and therapeutic approaches for neurodegenerative diseases. Prog. Retin. Eye Res. 43, 17–75 (2014).

74. Jones, B. W. et al. Retinal Remodeling and Metabolic Alterations in Human AMD. Front. Cell. Neurosci. 10, (2016).

75. Pandiyan, V. P., Jiang, X., Kuchenbecker, J. A. & Sabesan, R. Reflective mirror-based line-scan adaptive optics OCT for imaging retinal structure and function. Biomed. Opt. Express 12, 5865 (2021).

76. Pandiyan, V. P. et al. Characterizing cone spectral classification by optoretinography. Biomed. Opt. Express 13, 6574 (2022).

77. Teng, P. Caserel-an open source software for computer-aided segmentation of retinal layers in optical coherence tomography images. Zenodo (2013).

78. Stevenson, S. B. & Roorda, A. Correcting for miniature eye movements in high-resolution scanning laser ophthalmoscopy. in vol. 5688 145–151 (SPIE, 2005).

79. Scoles, D. et al. In Vivo Imaging of Human Cone Photoreceptor Inner Segments. Investig. Opthalmology Vis. Sci. 55, 4244 (2014).

