## Supplementary Information for "In vivo elastography of the human retina using light-evoked intrinsic actuation"

### 1. ORG signal extraction

We first removed the arbitrary phase offset at each pixel by referencing the phase of each volume to that of the first volume,

$$\tilde{I}'(x,y,z)^{(i)} = \tilde{I}(x,y,z)^{(i)} \cdot \exp\left[-j\arg\left(\tilde{I}(x,y,z)^{(1)}\right)\right], \tag{S1}$$

where $\tilde{I}(x,y,z)^{(i)}$ is the complex-valued OCT volume after registration. $x$, $y$ and $z$ denote the line, scan, and depth dimensions, respectively. $i$ is the volume index. $j$ represents the imaginary unit, and $\arg(\cdot)$ denotes the calculation of the argument.

#### 1.1 Axial-referencing analysis

In the axial-referencing analysis, the phase difference between two retinal bands was calculated to cancel common phase fluctuations. We first summed the time-referenced OCT signals from each band along the axial dimension,

$$\tilde{I}_{\text{ref}}(x,y)^{(i)} = \sum_{z=z_{r1}}^{z_{r2}} \tilde{I}'(x,y,z)^{(i)}, \tag{S2}$$

$$\tilde{I}_{\text{tar}}(x,y)^{(i)} = \sum_{z=z_{t1}}^{z_{t2}} \tilde{I}'(x,y,z)^{(i)}, \tag{S3}$$

where $z_{r1}$ and $z_{r2}$ denote the upper and lower boundaries of the reference band, while $z_{t1}$ and $z_{t2}$ represent the upper and lower boundaries of the target band.

The phase difference between these two retinal layers was then obtained by multiplying $\tilde{I}_{\text{tar}}(x,y)^{(i)}$ by the complex conjugate of $\tilde{I}_{\text{ref}}(x,y)^{(i)}$

$$\tilde{I}_{\text{axial}}(x,y)^{(i)} = \tilde{I}_{\text{tar}}(x,y)^{(i)} \tilde{I}^{*}_{\text{ref}}(x,y)^{(i)}, \tag{S4}$$

where $*$ denotes the complex conjugate operation.

To enhance the SNR of the phase trace, we applied a spatial kernel prior to phase extraction,

$$\phi_{\text{axial}}(x,y)^{(i)} = \arg\left(\tilde{I}_{\text{axial}}(x,y)^{(i)} \otimes g_1(x,y)\right), \tag{S5}$$

where $\otimes$ denotes the convolution operation. In this study, $g_1(x,y)$ was set to a $3 \times 3$ normalized boxcar kernel.

#### 1.2 Lateral-referencing analysis

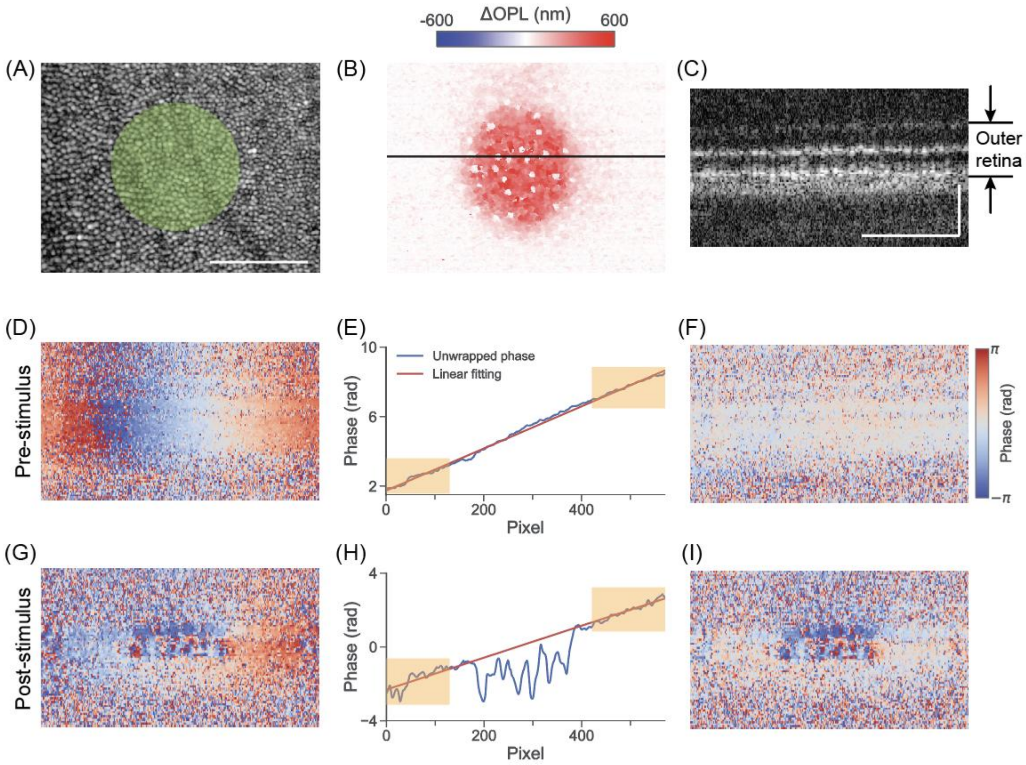

Figure S1. Illustration of the localized stimulus lateral-referencing analysis. (A) Structural *en-face* image at the cone outer segment tip (COST). Scale bar: 100 $\mu m$. (B) The conventional axial-referencing method revealed $\Delta$OPL along the cone outer segment (COS) in response to a localized visual stimulus (green spot in panel A). $\Delta$OPL was averaged over 0.4-0.6 s post-stimulus. (C) Cross-sectional structural image along the black line in panel B. Scale bar: 100 $\mu m$. (D) The phase difference between a pre-stimulus frame (same as panel C) and the corresponding frame in the first volume. (E) The phase difference map in panel D1 was axially averaged over the outer retina region (see panel C) and unwrapped along the line dimension (blue curve). The red line indicates a linear fit applied to regions outside the stimulus spot (see yellow rectangles). (F) Subtracting of the fitted linear phase trace (red line) from panel D yielded the lateral-referenced phase map. (G), (H) and (I) are the same as (D), (E) and (F), but show the phase difference between a post-stimulus frame and the corresponding frame in the first volume.

In the local-referencing analysis, a localized visual stimulus was delivered to the retina (green spot in Fig. S1A), and phase changes were calculated relative to the surrounding non-stimulated regions. Figure S1B shows the stimulus-evoked ΔOPL map along the cone outer segment (OS) obtained using the axial-referencing analysis, confirming that the stimulus was limited to a localized region. Figure S1D provides the phase difference between a pre-stimulus frame along the black line in Fig. S1B and the corresponding frame in the first volume. The associated phase fluctuations, which primarily arise from translational and rotational bulk movements [1,2], vary linearly along the line direction. Figure S1G shows the phase difference between a post-stimulus frame along the black line in Fig. S1B and the corresponding frame in the first volume, and contains both the ORG signal and phase fluctuations caused by residual retinal movement. To isolate the phase fluctuations associated with the bulk retinal movement, we axially summed the time-referenced signal $\tilde{I}'(x,y,z)^{(i)}$ across the outer retina (see Fig. S1C), applied a Gaussian convolution kernel $g_2(x)$ (kernel size: 7), and extracted its phase component,

$$\phi(x,y)^{(i)} = \arg\left(\sum_{z=z_1}^{z_2} \tilde{I}'(x,y,z)^{(i)} \otimes g_2(x)\right), \tag{S6}$$

where $z_1$ and $z_2$ are the depth indices corresponding to the upper and lower boundaries of the outer retina. For each cross-sectional scan, we selected regions outside the localized stimulation area, denoted as $x_r$, and performed a linear fit to estimate the phase fluctuations caused by retinal bulk movement. Specifically, we first unwrapped $\phi(x,y)^{(i)}$ along the line dimension to obtain $\phi^{\mathcal{U}}(x,y)^{(i)}$ (see blue lines in Figs. S1E and H). A linear fit was then computed between $x_r$ and $\phi^{\mathcal{U}}(x_r,y)^{(i)}$ using the least-squares criterion, yielding $\phi^{\mathcal{U}}(x_r,y)^{(i)} = p_0(y)^{(i)}x_r + p_1(y)^{(i)}$, with $p_0(y)^{(i)}$ and $p_1(y)^{(i)}$ representing the linear fitting coefficients (see red lines in Figs. S1E and H). The fitted linear phase fluctuation was then removed from the time-referenced OCT signal, yielding the lateral-referenced ORG signal (Figs. S1F and I),

$$\tilde{I}_{\text{lateral}}(x,y)^{(i)} = \tilde{I}'(x,y,z)^{(i)} \cdot \exp\left[-j\left(p_0(y)^{(i)}x + p_1(y)^{(i)}\right)\right]. \tag{S7}$$

The lateral-referenced ORG signal for each retinal band, spanning $z_{t1}$ to $z_{t2}$, was then obtained by,

$$\phi_{\text{lateral}}(x,y)^{(i)} = \arg\left(\sum_{z=z_{t1}}^{z_{t2}} \tilde{I}'(x,y,z)^{(i)} \otimes g_3(x,y)\right), \tag{S8}$$

where a Gaussian kernel $g_3(x,y)$ (kernel size: 3×3) was applied prior to phase extraction to enhance the SNR.

### 2. Analytical model and inverse relations

#### 2.1 Model assumptions and governing equations

We formulated a biomechanical model to explain light-evoked displacements of retinal layers. Specifically, the phototransduction cascade in cone photoreceptors leads to an increase in pressure and an elongation of the cone OS. After reaching a quasi-static phase (the post-fast-rising plateau of the elongation), the residual pressure difference between the inside and outside of the cone OS, denoted as $\Delta p$, is counterbalanced by elastic restoring forces exerted by the supracone space (SCS) on the COST side and by the inner retina on the ELM side. The displacement of the COST ($d_{COST}$) can be calculated as,

$$d_{COST} = \frac{\Delta p}{K_{A,SCS}}, \tag{S9}$$

where $K_{A,SCS}$ represents the area-normalized spring constant (i.e., spring constants per reference area) of the restoring forces exerted by the SCS. Hereafter, we refer to this area-normalized spring constant as areal spring constant for simplicity.

Assuming that the force applied at the ISOS junction is fully transmitted to the ELM, the pressure at the top of the inner segment (IS), denoted as $\Delta p_{MZ}$, is expressed as,

$$\Delta p_{MZ} = \frac{A_{OS}}{A_{MZ}} \Delta p, \tag{S10}$$

where $A_{OS}$ and $A_{MZ}$ are cross-sectional areas of the outer segment and myoid zone (MZ), respectively.

The displacements of ELM ($d_{ELM}$) can be calculated as,

$$d_{ELM} = \frac{\Delta p_{MZ}}{K_{A,IR}} = \frac{A_{OS}\Delta p}{A_{MZ}K_{A,IR}}, \tag{S11}$$

where $K_{A,IR}$ denotes the areal spring constant of the restoring forces exerted by the inner retina.

The compression of the IS ($\Delta L_{IS}$) is given by,

$$\Delta L_{IS} = A_{OS}\Delta p/k_{IS}, \tag{S12}$$

where $k_{IS}$ is the stiffness of the IS (see Supplementary Section 4.4 for details).

The displacement of the ISOS junction ($d_{ISOS}$) can be calculated as the sum of the ELM displacement and the IS compression, i.e.,

$$d_{ISOS} = d_{ELM} + \Delta L_{IS} = \Delta p\left(\frac{A_{OS}}{A_{MZ}K_{A,IR}} + \frac{A_{OS}}{k_{IS}}\right), \tag{S13}$$

and the elongation of the OS, denoted as $\Delta L_{OS}$, is given by,

$$\Delta L_{OS} = d_{COST} + d_{ISOS} = \Delta p\left(\frac{1}{K_{A,SCS}} + \frac{A_{OS}}{A_{MZ}K_{A,IR}} + \frac{A_{OS}}{k_{IS}}\right). \tag{S14}$$

Notably, all displacements and deformations in this linear elastic model depend on $\Delta p$ (See Supplementary Section 5). To forego the requirement to deduce a unique value for $\Delta p$, we defined two dimensionless metrics to characterize the fractional displacement of retinal layers:

$$f_{ISOS} = \frac{d_{ISOS}}{\Delta L_{OS}} \ , \tag{S15}$$

$$f_{ELM} = \frac{d_{ELM}}{d_{ISOS}} \ , \tag{S16}$$

where $f_{ISOS}$ represents the ISOS displacement normalized by the OS elongation, and $f_{ELM}$ denotes the ratio of the ELM displacement to the ISOS displacement. Substituting Eqs. (S11), (S13) and (S14) into Eqs. (S15)-(S16) yields,

$$K_{A,SCS} = \frac{f_{ISOS}f_{ELM}}{1-f_{ISOS}}\frac{A_{MZ}}{A_{OS}}K_{A,IR}, \tag{S17}$$

$$k_{IS} = \frac{f_{ELM}}{1-f_{ELM}}A_{MZ}K_{A,IR} = \frac{1-f_{ISOS}}{f_{ISOS}(1-f_{ELM})}A_{OS}K_{A,SCS}, \tag{S18}$$

Equations (S17) and (S18) establish the relationship between the mechanical properties of different retinal layers and their deformations. In Supplementary Section 3, we further derive $K_{A,IR}$, $K_{A,SCS}$, and $k_{IS}$ from Young's moduli and geometric parameters of the relevant retinal structure. In Supplementary Section 4, we estimate the Young's moduli of the SCS and inner segment by fitting the model predictions into experimental results.

#### 2.2 Experimental extraction of $f_{ISOS}$ and $f_{ELM}$

Experimentally, because the RPE can be treated as a static reference (see Fig. 2E), $d_{ISOS}$ and $d_{ELM}$ were obtained by computing ΔOPL between the ISOS and RPE, and between the ELM and RPE, respectively, using the axial-referencing analysis. As summarized in Section 3.2, $f_{ISOS}$ and $f_{ELM}$ in the foveal region ($0.25^{\circ}$ eccentricity) and the parafovea ($4^{\circ}$ eccentricity) were calculated based on ORG results acquired using *protocol 1* (see Methods in the main text). These data included measurements from 10 subjects at the highest stimulus strength (14.1 $\times 10^6$ photons/μm$^2$ on the retina), and from 4 subjects at the middle (8.8 $\times 10^6$ photons/μm$^2$) and low (4.4$\times 10^6$ photons/μm$^2$) stimulus strengths.

#### 2.3 Closed-form inverse relations between $K_{A,IR}$, $K_{A,SCS}$, and $k_{IS}$

According to Eq. (S17), the areal spring constant of the SCS in the fovea can be calculated as,

$$K_{A,SCS}^{0.25^{\circ}} = \frac{f_{ISOS}^{0.25^{\circ}} f_{ELM}^{0.25^{\circ}}}{1-f_{ISOS}^{0.25^{\circ}}} \frac{A_{MZ}^{0.25^{\circ}}}{A_{OS}^{0.25^{\circ}}} K_{A,IR}^{0.25^{\circ}}, \quad \text{(S19)}$$

and the areal spring constant of the inner retina in the parafovea is given by,

$$K_{A,IR}^{4^{\circ}} = \frac{1-f_{ISOS}^{4^{\circ}}}{f_{ISOS}^{4^{\circ}} f_{ELM}^{4^{\circ}}} \frac{A_{OS}^{4.0^{\circ}}}{A_{MZ}^{4.0^{\circ}}} K_{A,SCS}^{4^{\circ}}, \quad \text{(S20)}$$

Assuming that effective SCS support does not change substantially between the sampled eccentricities, i.e., $K_{A,SCS}^{4^{\circ}} \approx K_{A,SCS}^{0.25^{\circ}}$, and substituting Eq. (S19) into Eq. (S20) yields,

$$K_{A,IR}^{4^{\circ}} = \frac{1-f_{ISOS}^{4^{\circ}}}{1-f_{ISOS}^{0.25^{\circ}}} \frac{f_{ISOS}^{0.25^{\circ}} f_{ELM}^{0.25^{\circ}}}{f_{ISOS}^{4^{\circ}} f_{ELM}^{4^{\circ}}} \frac{A_{OS}^{4.0^{\circ}} A_{MZ}^{0.25^{\circ}}}{A_{OS}^{0.25^{\circ}} A_{MZ}^{4.0^{\circ}}} K_{A,IR}^{0.25^{\circ}}, \quad \text{(S21)}$$

Based on Eqs. (S18), the stiffness of the IS in the fovea and parafovea can be calculated as,

$$k_{IS}^{0.25^{\circ}} = \frac{f_{ELM}^{0.25^{\circ}}}{1-f_{ELM}^{0.5^{\circ}}} A_{MZ}^{0.25^{\circ}} K_{A,IR}, \quad \text{(S22)}$$

$$k_{IS}^{4^{\circ}} = \frac{1-f_{ISOS}^{4^{\circ}}}{f_{ISOS}^{4^{\circ}}\left(1-f_{ELM}^{4^{\circ}}\right)} A_{OS}^{4^{\circ}} K_{A,SCS}^{4^{\circ}} = \frac{1-f_{ISOS}^{4^{\circ}}}{f_{ISOS}^{4^{\circ}}\left(1-f_{ELM}^{4^{\circ}}\right)} A_{OS}^{4^{\circ}} K_{A,SCS}^{0.25^{\circ}}, \quad \text{(S23)}$$

where we assume the areal spring constant of the SCS is the same in the fovea and parafovea. Substituting Eq. (S19) into Eq. (S23) yields,

$$k_{IS}^{4^\circ} = \frac{1-f_{ISOS}^{4^\circ}}{f_{ISOS}^{4^\circ}\left(1-f_{ELM}^{4^\circ}\right)} \frac{f_{ISOS}^{0.25^\circ} f_{ELM}^{0.25^\circ}}{1-f_{ISOS}^{0.25^\circ}} \frac{A_{MZ}^{0.25^\circ}}{A_{OS}^{0.25^\circ}} A_{OS}^{4^\circ} K_{A,IR}^{0.25^\circ} \tag{S24}$$

### 3. Finite-element anchors and calibrations

#### 3.1 Foveal inner retina anchor $K_{A,IR}$

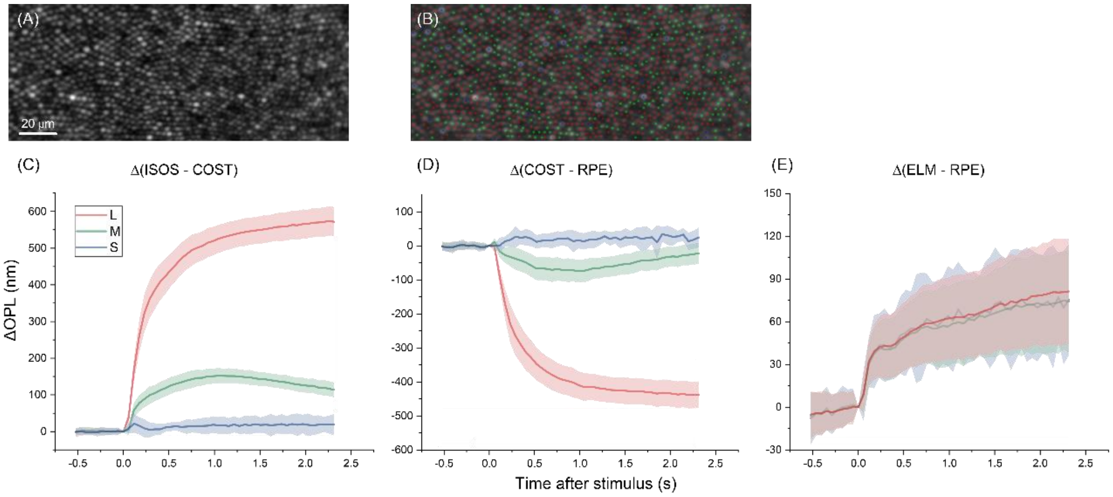

Figure S2. Various ORG signatures of individual cones measured using AO-OCT. (A) The maximum intensity projection (MIP) of the ISOS/COST interface. (B) Corresponding cone mosaic with different cone types obtained by ORG-based cone classification [3]. (C) Time-varying ORG traces of the cone outer segment (ISOS to COST) are shown for L-, M-, and S-cones. (D) Time-varying ORG traces spanning from COST to RPE for the L-, M- and S-cones. (E) Time-varying ORG traces spanning from ELM to RPE, where the ELM phase values are obtained from pixels directly overlying the inner segment of the L-, M- and S-cones shown in panel C&D. Shaded regions in panels (C)-(E) indicate ±1 standard deviation.

The ELM is a continuous adherens junction layer between the inner segments and the overlying Müller glial cells[4]. As shown in Fig. S2, while different cone types exhibit distinct responses at the COS following visual stimulation (see Fig. S2C&D), the ORG responses extracted between the ELM overlying the same cones and RPE are indistinguishable across spectral types (see Fig. S2E). This observation suggests that the ELM acts as an elastic membrane, and that forces originating from individual cones spread across the ELM interface. Accordingly, the pressure exerted on the ELM, denoted as $\Delta p_{ELM}$, is given by,

$$p_{ELM} = \alpha \Delta p_{MZ}, \tag{S25}$$

where $\alpha$ is the areal fraction occupied by the cone MZ and can be calculated as,

$$\alpha = \sigma_{\text{cone}} A_{MZ}, \tag{S26}$$

where $\sigma_{\text{cone}}$ is the areal cone density.

Here, we constructed a COMSOL model to calculate the areal spring constant of the ELM using parameters listed in Table S1. Figure S3 left provides the light-evoked ORG responses (Method *protocol 1*, $8.8 \times 10^6$ photons/μm$^2$) in the inner retina averaged across 50 adjacent B-scans and over 0.4-0.8 s post-stimulus. Notably, the inner retinal deformation is limited to the foveal region. This observation is consistent with the known biomechanical properties of the ILM. Specifically, the ILM is mechanically strong and 1000-fold stiffer than the cellular layers of the retina. More importantly, the ILM is very thin at the foveal center, and its thickness increases rapidly with retinal eccentricity over 300–1000 μm[5]. Accordingly, the ILM within 1 deg (300 μm) is set as a free boundary in the model, the corresponding inner retinal layers' thickness and biomaterial properties are shown in Fig. S3 and Table S1. Figure S3 right shows the COMSOL fitting results when the uniform pressure applied to ELM ($p_{ELM}$) was set to 1.2 Pa. The resulting ELM deformation in the foveal region was 75 nm, corresponding to a ΔOPL of 106 nm, assuming a refractive index of 1.41 [6,7]. In this region, the areal spring constant of the inner retina exerted to individual photoreceptors can be calculated as,

$$K_{A,IR}^{0.25^\circ} = \frac{\Delta p_{MZ}}{d_{ELM}} = \frac{1}{\alpha}\frac{p_{ELM}}{d_{ELM}} = 1.8 \times 10^7 \text{ N·m}^{-3} \tag{S27}$$

where $\alpha = 0.92$ was obtained using Eq. (S26), based on a cone density at 0.25° retinal eccentricity[8] of $1.5 \times 10^5$ cells/mm$^2$ and the corresponding MZ diameter of 2.8 μm [9].

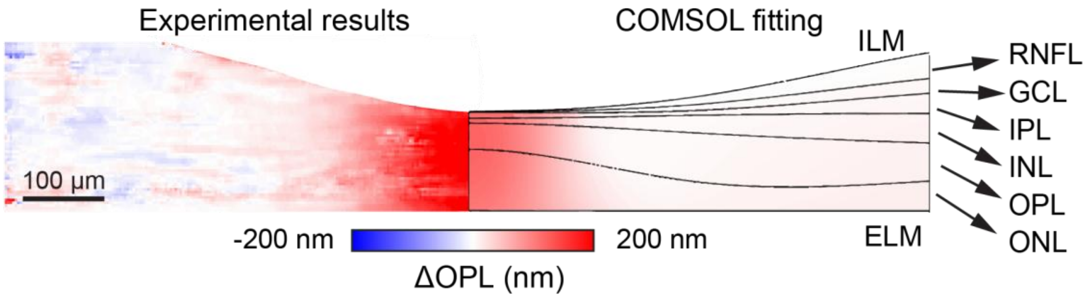

Figure S3. Comparison of stimulus-evoked ΔOPL measured in the inner retina and a COMSOL finite-element model. Left: experimentally measured ΔOPL map averaged across 50 adjacent B-scans and over 0.4-0.8 s post-stimulus. Right: COMSOL fit using a layered inner-retina configuration[10], with Young's modulus and Poisson's ratio of each layer provided in Table S1. The ILM within 1 deg (300 μm) was set as a free boundary, whereas the ILM beyond 1 deg was treated as a fixed boundary. The deformation predicted by the COMSOL model was converted into ΔOPL using a refractive index of 1.41 [6,7], and the uniform pressure applied to the ELM was adjusted so that the COMSOL model fitted experimental results. RNFL: retinal nerve fiber layer; GCL: ganglion cell layer; IPL: inner plexiform layer; INL: inner nuclear layer; OPL: outer plexiform layer; ONL: outer nuclear layer.

**Table S1. The Biomaterial properties of inner retinal layers**

| Inner retinal layers | Young's modulus | Poisson's ratio |
|---|---|---|
| Retinal Nerve Fiber Layer (RNFL) | 1.3 kPa[2,11] | 0.47 |
| Ganglion cell layer (GCL) | 1.2 kPa[2,11] | 0.47 |
| Inner Plexiform Layer (IPL) | 1.3 kPa[2,11] | 0.47 |
| Inner Nuclear Layer (INL) | 2.7 kPa[2,11] | 0.47 |
| Outer Plexiform Layer/Henle's Fiber Layer (OPL/HFL) | 8.0 kPa[2,11] | 0.47 |
| Outer Nuclear Layer (ONL) | 2.7 kPa[2,11] | 0.47 |

#### 3.2 SCS calibration from $K_{A,SCS}$ to $E_{SCS}$

As shown in Fig. S2D, the light-evoked displacements of different cone types are distinct at the COST level. Accordingly, we built a COMSOL simulation to compute the areal spring constant of the SCS at the individual cone level. As shown in Fig. S4A, a uniform pressure was applied to the end of the COS, which was modeled as a rigid cylinder with a diameter of 1.3 μm and supported by a 1-μm-thick SCS layer (Poisson's ratio: 0.45)[4,12]. The bottom of the SCS was fixed, consistent with experimental observations that the RPE serves as a static boundary (see Fig. 2B&2C). This linear elastic model shows that the areal spring constant of the SCS scales linearly with the Young's modulus of the SCS ($E_{SCS}$) as,

$$K_{A,SCS}^{0.25^\circ} = aE_{SCS}, \tag{S28}$$

where the coefficient $a = 3.2 \times 10^6$ m$^{-1}$. Moreover, the axial displacement (see Fig. S4B) shows that the COST displacement has a very limited spatial extent (FWHM = 1.64 μm), supporting our observations that COS responses from individual cones can be readily differentiated (see Fig. S2).

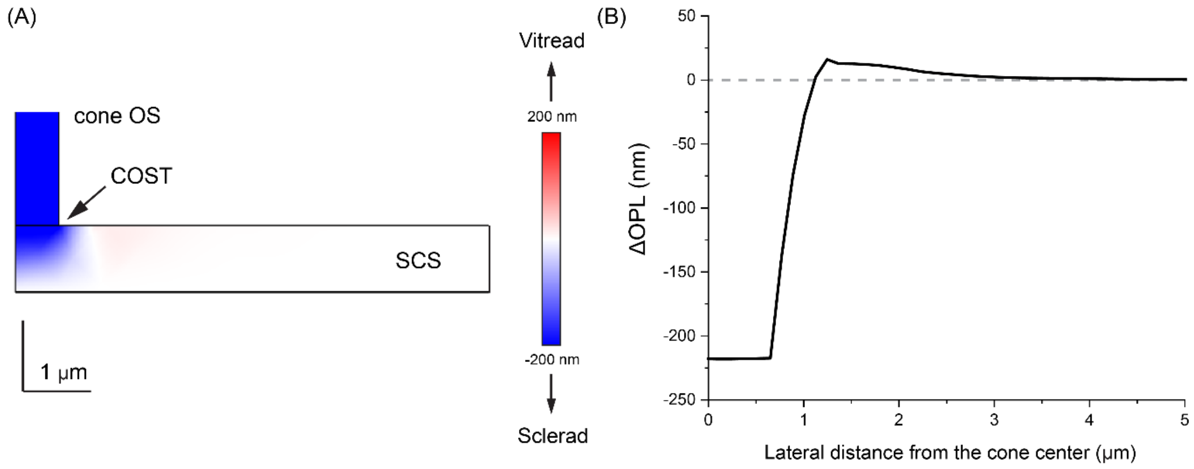

Figure S4. COMSOL modeling of a cone OS and its underlying SCS. A uniform pressure of 6.0 Pa was applied to COST. (A) The symmetrical half side of 2D axial displacement field. (B) The axial displacement at the depth of COST.

This section provides the conversion between the areal spring constant and Young's modulus of the SCS, with detailed values given in Supplementary Section 4.2.

#### 3.3 Full COMSOL forward validation of multilayer deformation

After parameter inference, the final parameter set was inserted into a full COMSOL model to visualize the quasi-static deformation field in foveal and parafoveal cones. The simulated ELM, ISOS, and COST motions reproduced the experimentally measured deformation partitioning and are presented here as a forward validation and visualization of the analytically inferred parameters, shown in Fig. S5.

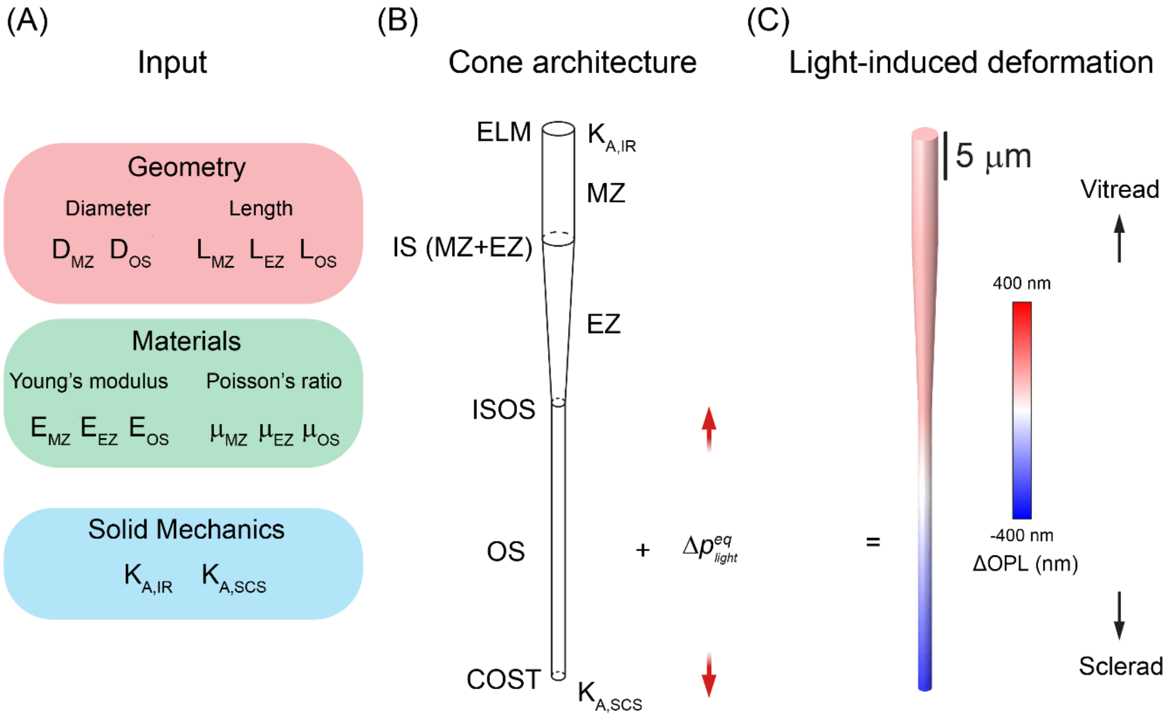

Figure S5. COMSOL workflow for simulating light-induced cone deformation. (A) Input parameters used to construct the finite element cone photoreceptor model include cone geometry, region-specific material properties, and solid mechanics boundary parameters. Geometry inputs include the MZ and OS diameters ($D_{MZ}$ , $D_{OS}$) and the lengths of the MZ, EZ, and OS ($L_{MZ}$, $L_{EZ}$, $L_{OS}$). Material inputs include Young's moduli ($E_{MZ}$, $E_{EZ}$, $E_{OS}$) and Poisson's ratios ($\mu_{MZ}$, $\mu_{EZ}$, $\mu_{OS}$), while mechanical coupling to the inner retina and supracone space is represented by areal spring constants $K_{A,IR}$ and $K_{A,SCS}$, respectively. (B) The inferred parameter set was incorporated into a full COMSOL solid mechanics model of a cone spanning the ELM to COST, with anatomical interfaces including the ISOS. A light-induced equilibrium pressure perturbation, $\Delta p^{eq}_{light}$, was applied to drive quasi-static deformation. (C) The resulting deformation field visualizes the predicted light-induced optical path length change ($\Delta \mathrm{OPL}$), with displacement toward the vitreous shown as positive and displacement toward the sclera shown as negative. The simulated ELM, ISOS, and COST motions reproduced the experimentally measured deformation partitioning, providing forward validation and visualization of the analytically inferred parameters in foveal and parafoveal cones.

### 4. Parameter estimation and sensitivity analysis

#### 4.1 Experimental fraction values used for parameter estimation

The experimental fractions of $f_{ISOS}$ and $f_{ELM}$ in the fovea and parafovea were calculated as described in Supplementary Section 2.2 and are summarized in Table S2 below.

**Table S2. The ORG fraction from experimental results**

| Parameter | Fovea | Parafovea |
|---|---|---|
| $f_{\mathrm{ISOS}}$ | 0.31 ± 0.05 | 0.19 ± 0.03 |
| $f_{\mathrm{ELM}}$ | 0.74 ± 0.04 | 0.08 ± 0.04 |

#### 4.2 Solving for the foveal parameters

Using the foveal $K_{A,\mathrm{IR}}$ obtained in Supplementary Section 3.1 together with the measured $f_{\mathrm{ISOS}}$ and $f_{\mathrm{ELM}}$ values in Table S2 and cone geometry provided in Table S3, Eqs. (S19) and (S22) yields $K_{A,SCS}^{0.25^\circ} = (2.8 \pm 0.7) \times 10^7$ N·m$^{-3}$ and $k_{IS}^{0.25^\circ} = (3.3 \pm 0.7) \times 10^{-4}$ N·m$^{-1}$. The obtained $K_{A,\mathrm{SCS}}$ can be further converted to $E_{\mathrm{SCS}}$ using the calibration in Supplementary Section 3.2, resulting in $E_{SCS}^{0.25^\circ} = 8.7 \pm 2.0$ Pa.

#### 4.3 Estimating the parafoveal anchors

With the measured $f_{\mathrm{ISOS}}$ and $f_{\mathrm{ELM}}$ in Table S2, Eqs. (S21) and (S24) yielded $K_{A,IR}^{4^\circ} = (5.3 \pm 2.3) \times 10^7$ N·m$^{-3}$and $k_{IS}^{4^\circ} = (1.6 \pm 0.3) \times 10^{-4}$ N·m$^{-1}$.

#### 4.4 EZ/MZ sensitivity analysis

To assess the compression of the IS, we treated the myoid zone (MZ) and ellipsoid zone (EZ) separately. Assuming that the force applied at the ISOS junction is fully transmitted to the ELM and is uniformly distributed across the IS at each depth, the compression of the MZ, modeled as a cylinder, can be calculated as,

$$\Delta L_{MZ} = \frac{4FL_{MZ}}{\pi E_{MZ} D_{MZ}^2}, \qquad \text{(S29)}$$

where $L_{MZ}$, $D_{MZ}$ and $E_{MZ}$ are the length, diameter, and Young's modulus of the MZ, respectively. The transmitted force $F$ is given by,

$$F = \Delta p \cdot A_{OS}, \tag{S30}$$

The EZ was modeled as a tapered frustum, and its cross-sectional diameter was defined as a function of the axial distance from the ISOS, denoted by $z$,

$$D_{EZ}(z) = D_{OS} + \frac{D_{MZ} - D_{OS}}{L_{EZ}} z, \text{ with } 0 \le z \le L_{EZ}, \tag{S31}$$

where $L_{EZ}$ is the length of the EZ.

The compression of the EZ can be computed by,

$$\Delta L_{EZ} = \int_0^{L_{EZ}} \frac{4F}{\pi E_{EZ} D_{EZ}^2(z)} dz, \tag{S32}$$

where $E_{EZ}$ denotes the Young's modulus of the EZ.

Substituting Eq. (S31) into Eq. (S32), $\Delta L_{EZ}$ can be expressed as,

$$\Delta L_{EZ} = \frac{4F}{\pi E_{EZ}} \cdot \frac{L_{EZ}}{D_{MZ} D_{OS}}. \tag{S33}$$

Accordingly, the compression of the IS ($\Delta L_{IS}$) is the sum of the compression of the MZ and EZ, i.e.,

$$\Delta L_{IS} = \Delta L_{MZ} + \Delta L_{EZ} = \left( \frac{4L_{MZ}}{\pi E_{MZ} D_{MZ}^2} + \frac{4L_{EZ}}{\pi E_{EZ} D_{MZ} D_{OS}} \right) F. \tag{S34}$$

The stiffness of the cone IS can then be written as,

$$k_{IS} = \frac{F}{\Delta L_{IS}} = \left( \frac{4L_{MZ}}{\pi E_{MZ} D_{MZ}^2} + \frac{4L_{EZ}}{\pi E_{EZ} D_{MZ} D_{OS}} \right)^{-1}, \tag{S35}$$

Since IS stiffness is determined by both the geometric parameters[4] (summarized in Table S3, the length of MZ and EZ is based on the ratio mentioned in ref[4] and applied into our measured IS length accordingly, i.e., MZ length + EZ length = IS length) and the Young's moduli of the EZ and MZ. For a given $k_{IS}$ and $E_{EZ}$, $E_{MZ}$ can be computed as,

$$E_{MZ} = \frac{4L_{MZ}}{\pi D_{MZ}^2} \left( \frac{1}{k_{IS}} - \frac{4L_{EZ}}{\pi E_{EZ} D_{MZ} D_{OS}} \right)^{-1}, \tag{S36}$$

Here, we performed a sensitivity analysis by sweeping $E_{EZ}$ over a plausible range (10-200 kPa). For each assumed $E_{\mathrm{EZ}}$, we substituted the value of $k_{IS}$ calculated from each measurement (Eq. S22 and Eq. S24) into Eq. (S36) and finally obtained the mean value and uncertainty of $E_{MZ}$ in the fovea and parafovea (see Fig. S6). We found that the value of $E_{MZ}$ is relatively robust to the choice of $E_{EZ}$. When $E_{EZ}$ was fixed at $40\,\mathrm{kPa}$, $E_{MZ}^{0.25^\circ} = 0.67 \pm 0.14\,\mathrm{kPa}$ and $E_{MZ}^{4^\circ} = 0.05 \pm 0.01\,\mathrm{kPa}$.

**Table S3. Pre-defined cone geometric parameter**

| Parameter | Fovea | Parafovea | Units |
|---|---|---|---|
| MZ length, $L_{\mathrm{MZ}}$ | 12 | 10 | μm |
| EZ length, $L_{\mathrm{EZ}}$ | 18 | 12 | μm |
| Outer-segment length, $L_{\mathrm{OS}}$ | 31 | 20 | μm |
| Myoid zone diameter*, $D_{MZ}$ | 2.8[9] | 6.5[9] | μm |
| Outer-segment diameter, $D_{\mathrm{OS}}$ | 1.3[12] | 1.3[12] | μm |

*: In the model, the photoreceptor IS was represented as a cylindrical MZ and a tapering frustum-shaped EZ. Because the myoid and ellipsoid are contiguous subregions of the IS, we imposed radial continuity at the MZ–EZ interface and set the MZ diameter equal to the IS diameter (proximal EZ diameter[8]).

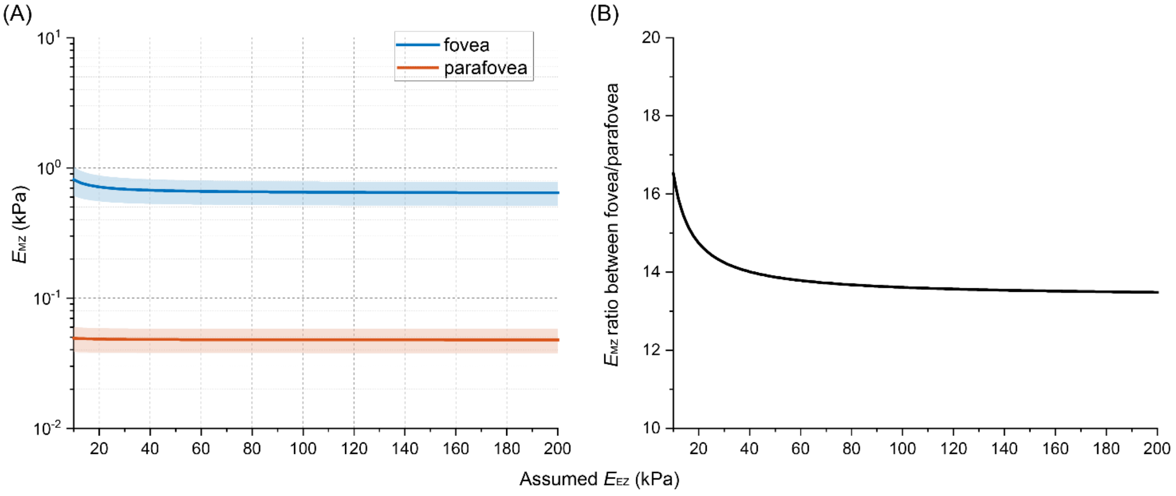

Figure S6. Inner-segment decomposition: (A) inferred MZ modulus as a function of assumed EZ modulus of range between 10 – 200 kPa. (B) The ratio of MZ modulus with eccentricity-dependent (fovea/parafovea) when assumed EZ modulus varies.

#### 4.5 Summary of inferred parameters

The biomechanical parameters of the retinal structures predicted by our model (see Supplementary Section 2) are summarized in Table S4.

**Table S4. Inferred parameters between fovea and parafovea**

| Quantity | Fovea | Parafovea* | Typical value** | Unit |
|---|---|---|---|---|
| Areal spring constant of the inner retina, $K_{A,IR}$ | $1.8 \times 10^7$ | $(5.3 \pm 2.3) \times 10^7$ | $3.0 - 6.0 \times 10^7$ (parafovea) | N/m³ |
| Areal spring constant of the SCS, $K_{\mathrm{A,SCS}}$ | $(2.8 \pm 0.6) \times 10^7$ | $(2.8 \pm 0.6) \times 10^7$ | / | N/m³ |
| Young's modulus of the SCS, $E_{SCS}$ | $8.7 \pm 2.0$ | / | $10 - 100$ | Pa |
| IS stiffness, $K_{\mathrm{IS}}$ | $(3.3 \pm 0.7) \times 10^{-4}$ | $(1.6 \pm 0.3) \times 10^{-4}$ | / | N/m |
| Young's modulus of the EZ, $E_{EZ}$*** | 40 | 40 | $10 - 115$ | kPa |

| | | | | |
|---|---|---|---|---|
| Young's modulus of the MZ, $E_{MZ}$ | $0.67 \pm 0.14$ | $0.05 \pm 0.01$ | $0.03 - 2.0$ | kPa |

*: Absolute values in the parafovea were derived based on the assumption that the areal spring constant of the SCS does not change substantially between the fovea and parafovea.

**: Typical values for $K_{A,IR}$ of parafovea estimated using $K_A \approx E/h$, with human macular and inner-retinal thicknesses from OCT studies[13,14] and apparent Young's modulus of the living inner retina from scanning-force microscopy[15]. The $E_{SCS}$ value was treated as an effective supracone/interphotoreceptor-matrix modulus; direct human SCS mechanical measurements were not found, so the estimate was based on the interphotoreceptor matrix being a hydrated extracellular matrix surrounding photoreceptors and occupying the subretinal/supracone region[16,17], together with reported low-stiffness hyaluronan-based hydrogel mechanics[18,19]. The $E_{EZ}$ value was assigned from the anatomical identification of the EZ with the mitochondria-rich photoreceptor ellipsoid[20]. The $E_{MZ}$ value was treated as an effective soft myoid/cytoplasmic modulus for the photoreceptor myoid zone; because direct human MZ measurements were not available, the range was estimated from retinal/CNS cell viscoelastic measurements[21], living-neuron AFM moduli[22], and the anatomical identification of the MZ as the photoreceptor inner-segment myoid between the mitochondria-rich EZ and the photoreceptor nucleus[4].

***: $E_{EZ}$ and $E_{MZ}$ cannot be obtained independently in this study. Given that a significant fraction of EZ is constituted by mitochondria[23], we assume $E_{EZ}$ = 40 kPa. A detailed analysis of $E_{MZ}$ values obtained by varying $E_{EZ}$ value is provided in Fig. S6.

### 5. Linking the light-induced OS pressure increase and deformation

#### 5.1 Total equilibrium OS pressure versus change in OS length

To estimate the total equilibrium pressure generated within the OS at the quasi-static state–including the pressure $\Delta p$ transmitted to the inner retina and SCS, as well as the component required to counterbalance the OS restoring force itself–we used a forward calibration approach described in Supplementary Section 3.3, with the OS Young's modulus fixed at 26 kPa [11]. The OS elongation (quantified here as ΔOPL using a refractive index of 1.41) scales linearly with the input total equilibrium pressure over 60.9–609 Pa, yielding an OS elongation range of the 100–1000 nm range and a slope of 1.64 nm/Pa for foveal cones (Fig. S7).

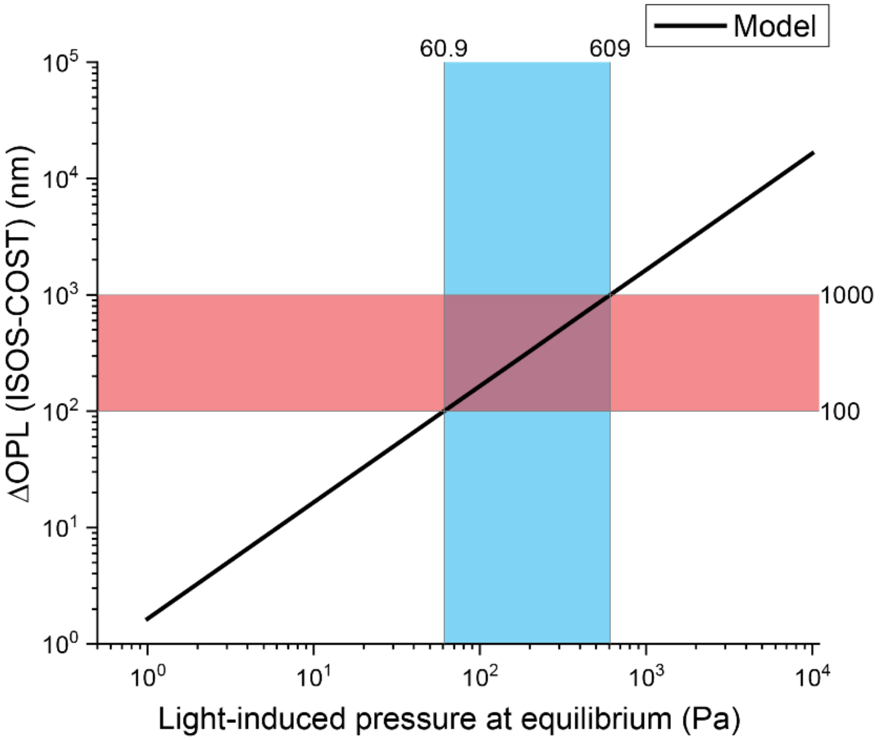

Figure S7. Forward calibration between total equilibrium OS pressure and OS elongation**.** Note that the x-axis represents the total equilibrium pressure required to produce the specified OS elongation and therefore includes both the residual transmitted pressure $\Delta p$ used in the analytical spring series model and the pressure needed to counterbalance the OS restoring force itself.

#### 5.2 Distinction between total OS pressure and residual transmitted pressure Δ*p*

The pressure reported in Fig. S7 is the total equilibrium pressure generated within the OS that is required to produce the measured OS elongation. A portion of this total equilibrium pressure is expended in stretching the OS itself, whereas the residual component ($\Delta p$) serves as the input for the analytical spring-series model and is transmitted to the proximal ELM/inner-retina side and the distal SCS side. The total light-induced pressure at the quasi-static state, denoted as $\Delta p_{light}^{eq}$, is related to the residual transmitted pressure ($\Delta p$) by,

$$\Delta p_{light}^{eq} = \frac{k_{os}\Delta L_{OS}}{A_{os}} + \Delta p, \tag{S37}$$

where $k_{os}$ is the spring constant of the cone OS given by $k_{os} = E_{OS}\,A_{OS}/L_{OS}$. Thus, Eq. (S37) can be rewritten as,

$$\Delta p_{light}^{eq} = \frac{\Delta L_{OS}}{L_{OS}} E_{os} + \Delta p, \tag{S38}$$

Note that although the value of $\Delta p_{light}^{eq}$ depends on the OS Young's modulus ($E_{os}$), variations in $E_{os}$ do not affect the retinal deformation derived from $\Delta p$ and, accordingly, do not affect the relative stiffness of retinal structures.

Combining Eq. (S9) with the definition of $f_{ISOS}$, the ratio between OS elongation and the total equilibrium pressure can be written as:

$$\frac{\Delta L_{OS}}{\Delta p_{light}^{eq}} = \left[\frac{E_{os}}{L_{os}} + K_{A,SCS}(1 - f_{ISOS})\right]^{-1} \approx \frac{L_{os}}{E_{os}}, \tag{S39}$$

where the approximation is valid because $E_{os}/L_{OS}$ is approximately 40- and 60-fold larger than $K_{A,SCS}(1 - f_{ISOS})$ for foveal and parafoveal cones, respectively. This indicates that a large fraction of the total equilibrium pressure is used to counterbalance the restoring force exerted by the OS itself. According to Eq. (S39), assuming the same OS Young's modulus for foveal and parafoveal cone outer segments, the ratio between OS elongation, quantified by ΔOPL, and total equilibrium pressure for parafoveal cones can be obtained by scaling the foveal value (Supplementary Section 5.1) by the ratio of their OS lengths (Table S3) as: $1.64\ \mathrm{nm/Pa} \times (20\ \mu\mathrm{m}/31\ \mu\mathrm{m}) =\ 1.06\ \mathrm{nm/Pa}$.

### Supplementary Figures

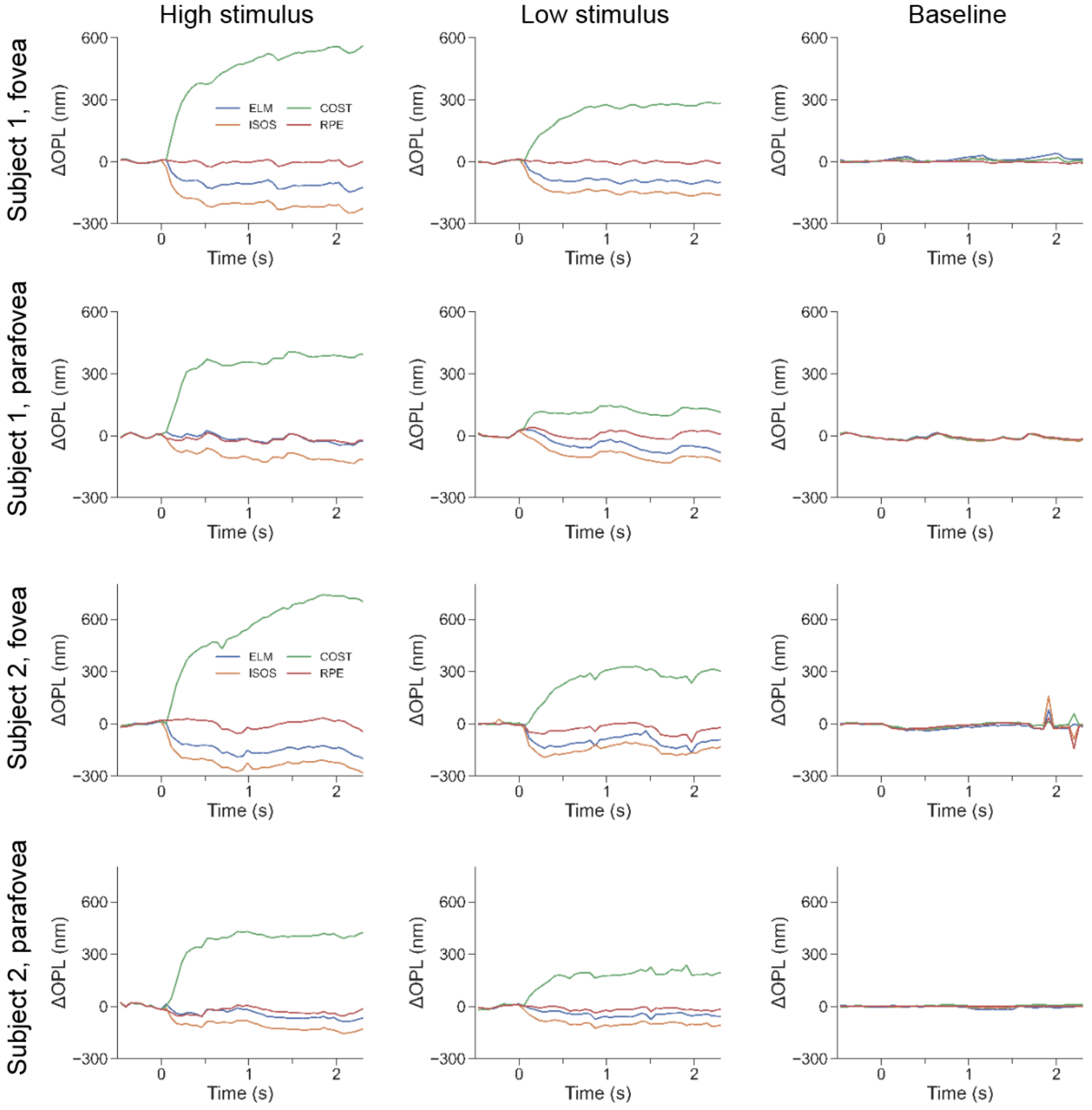

Figure S8. Local-referenced ORG signals measured from two subjects at two retinal eccentricities (fovea and parafovea). The first and second columns show ORG signals in response to high ($8.8\times10^{6}$ photons/μm$^{2}$) and low ($2.2\times10^{6}$ photons/μm$^{2}$) stimulus strengths, respectively. The last column shows signals recorded without visual stimulation.

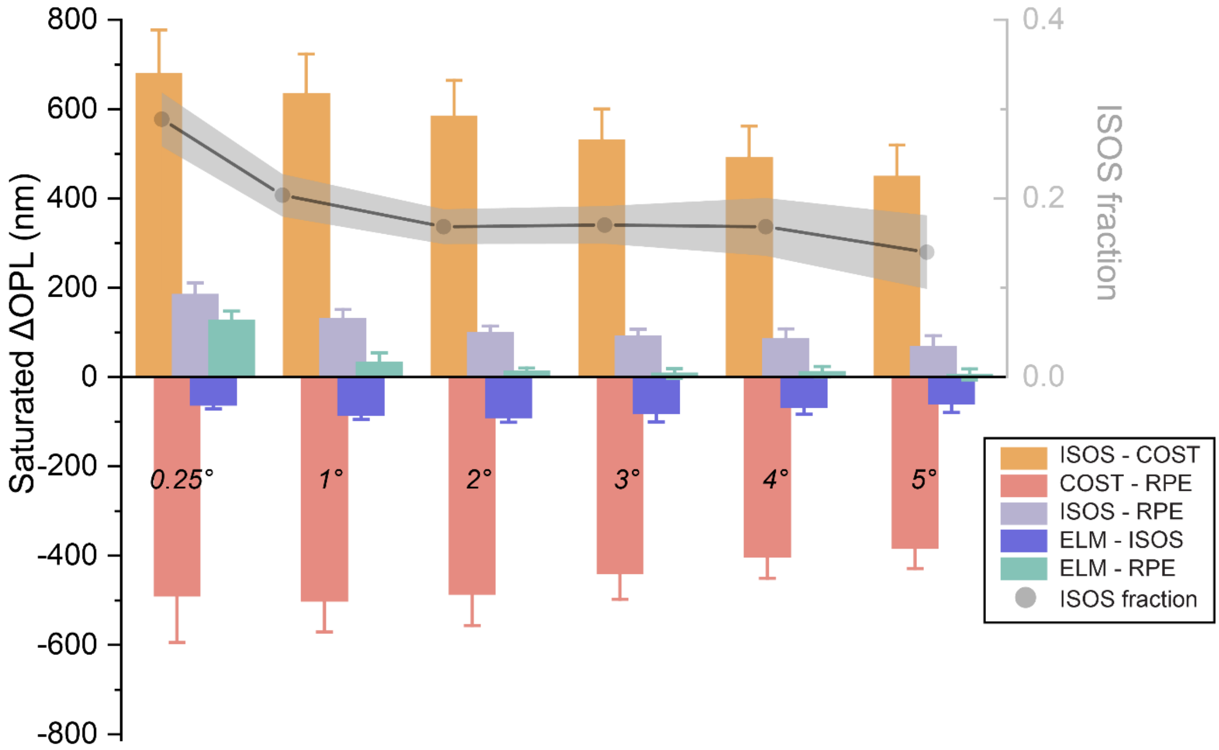

Figure S9: Eccentricity-dependence of the layer deformations. The light-evoked ΔOPL was sampled from the fovea to the parafovea (as shown Fig. 3A) and the post-fast-rising (t = 0.4 – 0.6 s) ΔOPL amplitude at stimulus strength of $14.1\times10^6$ photons/$\mu m^2$ is plotted as a function of eccentricity in an expanded cohort (n = 10 subjects). Overall, the magnitude of ISOS–COST exhibited the largest amplitude and decreased monotonically with eccentricity, indicating that stimulus-evoked OS expansion is strongest near the fovea. Because the RPE served as a static reference layer, COST–RPE and ISOS–RPE reflected the shifts of COST and ISOS interfaces, respectively and both amplitudes decreased with increasing eccentricity. The relative contributions of COST and ISOS to the COS elongation can be distinguished by the ISOS fraction (right y-axis): near the fovea, the vitread ISOS component accounted for approximately 25–30% of the total OS elongation (ISOS–COST), whereas in the parafovea, this fraction reduced to ~15–20%.

### Supplementary Table

**Table S5. Characteristics of clinical participants**

| Patient ID | Age Range | Sex | Genotype | Presenting Symptoms |
|---|---|---|---|---|
| RP01 | 26 – 30 | F | Indeterminate | nyctalopia, myopia, cystoid macular edema |
| RP02 | 26 – 30 | M | *IMPDH1*<br>c.942_944del, p.(Lys314del) | reduced peripheral vision, impaired dark adaptation |
| RP03 | 56 – 60 | F | *RHO*<br>c.165C>A, p.(Asn55Lys) | nyctalopia, increased glare while driving |
| RP04 | 21 – 25 | F | *PRPF31*<br>c.323-1G>T | nyctalopia, high myopia, blurring of central vision with flashes |
| RP05 | 36 – 40 | F | *USH2A*<br>*c.1558T>A, p.(Cys520Ser);*<br>*c.12575G>A, p.(Arg4192His)* | nyctalopia, central flashes |

**Supplementary References**

1. Veysset, D. *et al.* Interferometric imaging of thermal expansion for temperature control in retinal laser therapy. *Biomed. Opt. Express* **13**, 728 (2022).
2. Zhuo, Y. *et al.* Retinal thermal deformations measured with phase-sensitive optical coherence tomography in vivo. *Light Sci. Appl.* **14**, 151 (2025).
3. Pandiyan, V. P. *et al.* Characterizing cone spectral classification by optoretinography. *Biomed. Opt. Express* **13**, 6574 (2022).
4. Spaide, R. F. & Curcio, C. A. ANATOMICAL CORRELATES TO THE BANDS SEEN IN THE OUTER RETINA BY OPTICAL COHERENCE TOMOGRAPHY: Literature Review and Model. *Retina* **31**, 1609–1619 (2011).
5. Henrich, P. B. *et al.* Nanoscale Topographic and Biomechanical Studies of the Human Internal Limiting Membrane. *Investig. Opthalmology Vis. Sci.* **53**, 2561 (2012).
6. Sidman, R. L. The structure and concentration of solids in photoreceptor cells studied by refractometry and interference microscopy. *J. Biophys. Biochem. Cytol.* **3**, 15 (1957).
7. Pandiyan, V. P., Nguyen, P. T., Pugh, E. N. & Sabesan, R. Human cone elongation responses can be explained by photoactivated cone opsin and membrane swelling and osmotic response to phosphate produced by RGS9-catalyzed GTPase. *Proc. Natl. Acad. Sci.* **119**, e2202485119 (2022).
8. Curcio, C. A., Sloan, K. R., Kalina, R. E. & Hendrickson, A. E. Human photoreceptor topography. *J. Comp. Neurol.* **292**, 497–523 (1990).
9. Scoles, D. *et al.* In Vivo Imaging of Human Cone Photoreceptor Inner Segments. *Investig. Opthalmology Vis. Sci.* **55**, 4244 (2014).

10. Curcio, C. A. *et al.* Human Chorioretinal Layer Thicknesses Measured in Macula-wide, High-Resolution Histologic Sections. *Investig. Opthalmology Vis. Sci.* **52**, 3943 (2011).

11. Qu, Y. *et al.* Quantified elasticity mapping of retinal layers using synchronized acoustic radiation force optical coherence elastography. *Biomed. Opt. Express* **9**, 4054 (2018).

12. Yuodelis, C. & Hendrickson, A. A QUALITATIVE AND QUANTITATIVE ANALYSIS OF THE HUMAN FOVEA DURING DEVELOPMENT.

13. Arepalli, S. *et al.* Assessment of inner and outer retinal layer metrics on the Cirrus HD-OCT Platform in normal eyes. *PLoS One* **13**, e0203324 (2018).

14. Nieves-Moreno, M. *et al.* Normative database for separate inner retinal layers thickness using spectral domain optical coherence tomography in Caucasian population. *PLoS One* **12**, e0180450 (2017).

15. Franze, K. *et al.* Spatial mapping of the mechanical properties of the living retina using scanning force microscopy. *Soft Matter* **7**, 3147 (2011).

16. Ishikawa, M., Sawada, Y. & Yoshitomi, T. Structure and function of the interphotoreceptor matrix surrounding retinal photoreceptor cells. *Exp. Eye Res.* **133**, 3–18 (2015).

17. Hollyfield, J. G. Hyaluronan and the functional organization of the interphotoreceptor matrix. *Invest. Ophthalmol. Vis. Sci.* **40**, 2767–2769 (1999).

18. Xu, X., Jha, A. K., Harrington, D. A., Farach-Carson, M. C. & Jia, X. Hyaluronic acid-based hydrogels: from a natural polysaccharide to complex networks. *Soft Matter* **8**, 3280–3294 (2012).

19. Borzacchiello, A., Russo, L., Malle, B. M., Schwach-Abdellaoui, K. & Ambrosio, L. Hyaluronic acid based hydrogels for regenerative medicine applications. *BioMed Res. Int.* **2015**, 871218 (2015).

20. Li, Y., Honda, S., Iwami, K., Ohta, Y. & Umeda, N. Analysis of mitochondrial mechanical dynamics using a confocal fluorescence microscope with a bent optical fibre. *J. Microsc.* **260**, 140–151 (2015).

21. Lu, Y.-B. *et al.* Viscoelastic properties of individual glial cells and neurons in the CNS. *Proc. Natl. Acad. Sci.* **103**, 17759–17764 (2006).

22. Spedden, E., White, J. D., Naumova, E. N., Kaplan, D. L. & Staii, C. Elasticity Maps of Living Neurons Measured by Combined Fluorescence and Atomic Force Microscopy. *Biophys. J.* **103**, 868–877 (2012).

23. Hoang, Q. V., Linsenmeier, R. A., Chung, C. K. & Curcio, C. A. Photoreceptor inner segments in monkey and human retina: Mitochondrial density, optics, and regional variation. *Vis. Neurosci.* **19**, 395–407 (2002).
